# RoWS Biofabrication of Neuromuscular Tubuloids from Human Neuromesodermal Progenitors

**DOI:** 10.64898/2026.11.22.768485

**Authors:** Anita Florkowska, Shakeel Ahmad, Fabio Maiullari, Maryia Piletskaya, Maede Pajouhande, Saeed Rafieyan, Alicja Swiac, Marco Costantini

**Affiliations:** Institute of Physical Chemistry, Polish Academy of Sciences, Warsaw, Poland; Warsaw University of Technology, Chemistry Department, Warsaw, Poland

**Keywords:** Rotary wet-spinning, Neuromesodermal progenitors, Neuromuscular tubuloids, Core–shell hydrogels, Biofabrication, Tissue self-organization

## Abstract

Combining multicellular self-organization with defined tissue architecture remains a challenge in human neuromuscular modeling. Here, we use rotary wet-spinning (RoWS) biofabrication to encapsulate human neuromesodermal progenitors (NMPs) within fibrin-based cores surrounded by alginate shells, generating neuromuscular tubuloids (nmTubs). NMPs derived from a human embryonic stem cell and an induced pluripotent stem cell line were cultured for up to 50 days and assessed by gene expression analysis, immunofluorescence, tissue orientation measurements, and calcium imaging. Both cell lines generated muscle and neural compartments, with distinct differentiation kinetics and tissue proportions. Embryonic stem cell-derived nmTubs developed more extensive neural- and endothelial-associated structures and displayed spontaneous contractions from day 14 onward. Between days 30 and 50, muscle tissue area and myotube diameter increased, while acetylcholine receptor domains underwent spatial remodeling. Neural projections exhibited pronounced longitudinal organization within the constructs. Acetylcholine-associated calcium transients demonstrated tissue responsiveness, although functional neuromuscular transmission remains to be established. These findings show that RoWS can combine a common developmental progenitor population with a defined fibrous architecture to support multicellular neuromuscular development. nmTubs provide a platform to investigate the relationships among biomaterial geometry, tissue organization, and maturation in human neuromuscular models.

## 1. Introduction

Skeletal muscle development depends on coordinated interactions among cell populations whose migration, differentiation, and spatial organization give rise to functional tissue. Reproducing these processes *in vitro* remains challenging, particularly when muscle formation must be coupled with the development of a neuronal compartment. Although existing models generate differentiated myofibers, reconstructing the cellular interactions and spatial cues that underpin neuromuscular organization requires a shared three-dimensional environment that supports the development of both muscle and motor neurons [1,2].

Existing approaches address different aspects of this challenge. Two-dimensional differentiation systems provide experimental accessibility and control but lack the three-dimensional architecture of native tissue [3]. Neuromuscular organoids enable multicellular self-organization, although their geometry offers limited control over tissue orientation and the spatial arrangement of muscle and neural compartments [4,5]. Muscle-on-chip platforms provide greater control over microenvironmental conditions, but reproducing the coordinated emergence of both lineages during early development remains challenging [2,6]. Together, these approaches highlight the need for strategies that combine developmental self-organization with defined architectural guidance [7].

Our group previously developed rotary wet-spinning (RoWS) biofabrication to generate aligned, cell-laden hydrogel fibers for skeletal muscle and myotendinous junction engineering [8,9,10,11,12]. The anisotropic fiber environment guides myogenic organization and supports the formation of engineered muscle constructs with regenerative potential. However, our previous models lacked an intrinsic neuronal compartment and had limited satellite cell populations, thereby restricting their capacity to recapitulate the cellular diversity of neuromuscular development and regeneration. Incorporating progenitors capable of generating both neural and myogenic lineages could extend this platform toward more complex developmental tissue models.

Human neuromesodermal progenitors (NMPs) provide a developmental basis for this strategy. Derived from human pluripotent stem cells (hPSCs), these bipotent progenitors can generate neuroectoderm and paraxial mesodermal lineages, including motor neurons and skeletal muscle. This shared developmental origin provides a route to multicellular neuromuscular constructs through differentiation of a common progenitor population. Although this potential has been demonstrated in self-organizing neuromuscular models [4,13,14,15], the behavior of NMPs within geometrically confined hydrogel fibers remains unexplored.

Here, we combine RoWS biofabrication with human NMPs to generate elongated multicellular constructs termed neuromuscular tubuloids (nmTubs). NMPs are encapsulated within core–shell hydrogel fibers, where they proliferate, differentiate, and organize into muscle and neural compartments within a defined tubular geometry. Using NMPs derived from one human embryonic stem cell line (hESCs) and one induced pluripotent stem cell line (hiPSCs), we examine differences in differentiation trajectories, tissue composition, and structural maturation. The resulting nmTubs develop myofibers and neuronal networks, with neuronal projections closely associated with muscle regions containing acetylcholine receptor clusters. Spontaneous contractile activity varies between the two cell lines, while endothelial-associated populations and capillary-like structures indicate the emergence of additional cellular components.

Our findings establish nmTubs as a platform that combines the self-organizing potential of human NMPs with a biofabricated architecture for neuromuscular tissue development. Compared with organoid approaches that provide limited external control over tissue geometry, nmTubs introduce a defined longitudinal axis while preserving the capacity for multicellular self-organization (**Fig. 1**). Within this environment, developing tissues exhibit progressive structural organization, early spontaneous contractile activity (starting at day 14), and clustering of acetylcholine receptors in muscle regions associated with neuronal processes. This combination of developmental complexity and geometric control provides a basis for investigating the relationship between biomaterial architecture and human neuromuscular development, with potential applications in disease modeling and regenerative research.

**Figure 1.**
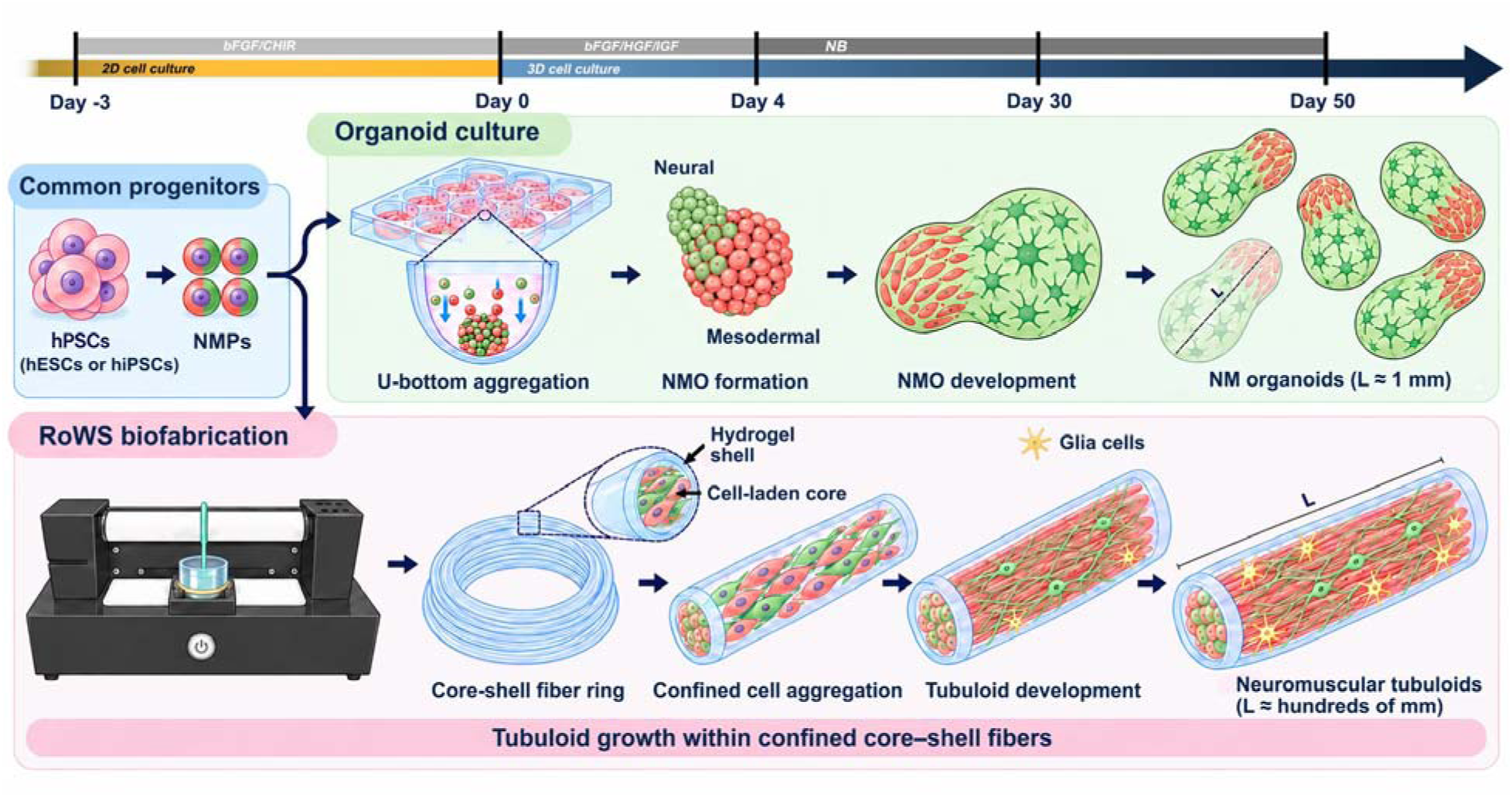
Study workflow and architectural differences between neuromuscular organoids and tubuloids. hPSCs are induced to differentiate into NMPs over three days using bFGF and CHIR99021. The upper branch illustrates conventional U-bottom aggregation and self-organization into neuromuscular organoids as a conceptual reference; the lower branch depicts the RoWS-based nmTub strategy investigated in this study. NMP-laden core–shell fibers provide a cell-supportive fibrin core and an alginate boundary that guides growth within an anisotropic geometry. nmTubs were cultured in Neurobasal-based medium (NB) supplemented with bFGF, HGF, and IGF1 during the first four days, followed by culture in NB through day 50. Red and green denote mesodermal/myogenic and neuroectodermal/neural populations, respectively; yellow denotes glial cells. Approximate lengths highlight the difference between millimeter-scale organoids and continuous fibers spanning tens of centimeters.

## 2. Results

### 2.1. Generation and self*-*organization *of* human neuromuscular *tubuloids*

To generate nmTubs, we first differentiated hPSCs into NMPs, a transient, bipotent developmental population capable of giving rise to spinal motor neurons and skeletal muscle, but not endodermal derivatives [16,17]. We used two hPSC lines: hESCs (H9 line) and hiPSCs (IIMCBi002-A line) [18]. Combined activation of WNT signaling and stimulation with fibroblast growth factor (FGF) for three days induced NMPs formation in both lines (**Fig. 2A**), following an established differentiation strategy [4,13,14]. To assess induction, we examined the characteristic NMP markers SOX2, T/BRACHYURY (T/BRACH), and CDX2 by gene expression analysis and evaluated SOX2 and T/BRACH co-expression by immunofluorescence (**Fig. 2B, C**). *T/BRACH* and *CDX2* mRNA levels increased significantly, consistent with the acquisition of nascent mesodermal characteristics and posterior axial identity, respectively [14,19,20]. *SOX2* mRNA expression did not increase significantly following induction (**Fig. 2B**); nevertheless, immunofluorescence revealed substantial overlap between SOX2 and T/BRACH signals in both hESC- and hiPSC-derived NMPs (**Fig. 2C**). Pearson’s correlation coefficients were 0.69 and 0.88, respectively, supporting co-expression of these defining NMP markers (**Fig. 2D**). This transition was accompanied by a significant reduction in *POU5F1* and *NANOG* expression relative to undifferentiated hPSCs, consistent with exit from pluripotency and acquisition of an NMP-associated phenotype (**Fig. S1A, B**).

**Figure 2.**
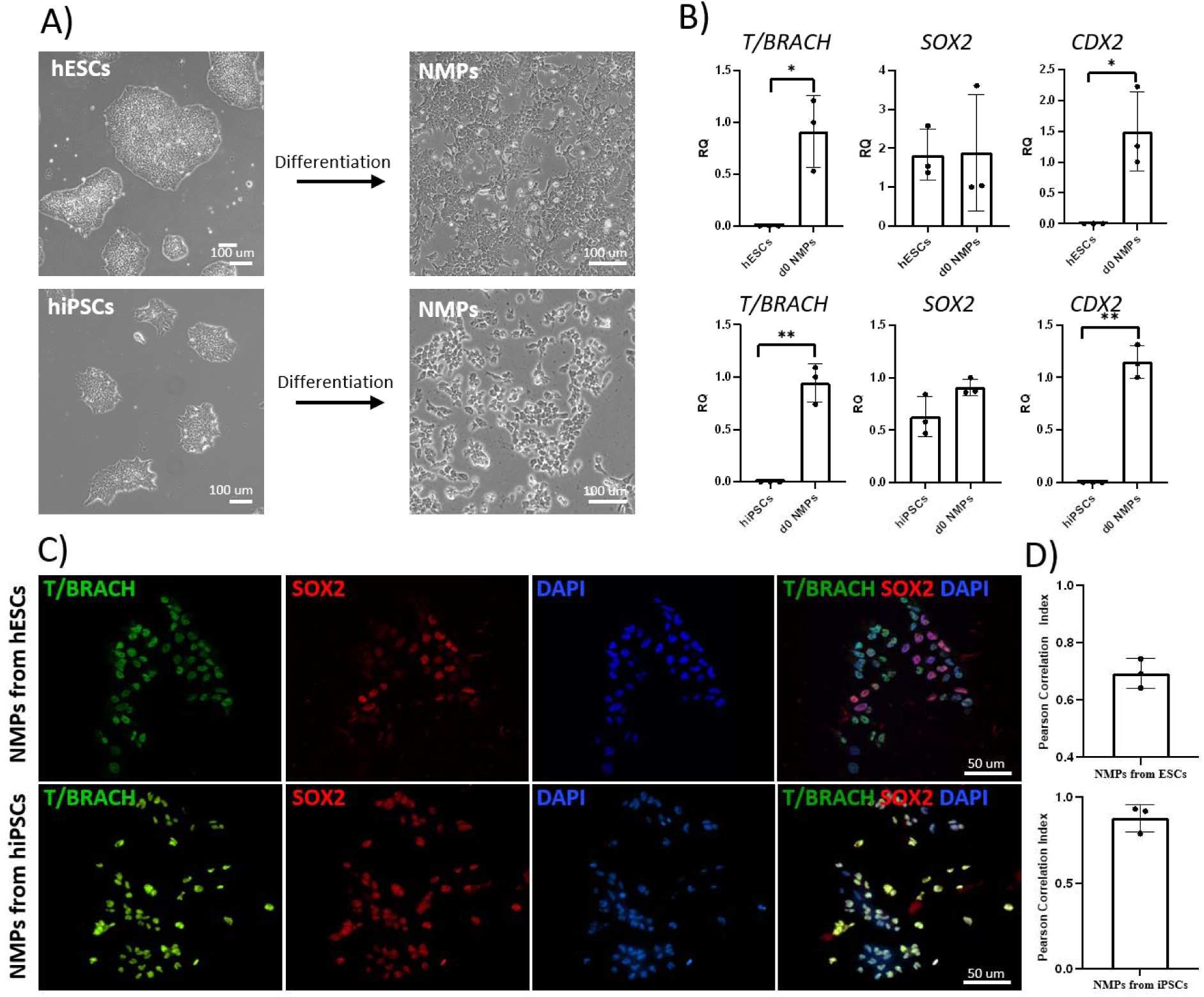
Induction of neuromesodermal progenitors from human pluripotent stem cells. **A)** Representative brightfield images of undifferentiated hESCs and hiPSCs and their NMP derivatives. Scale bar: 100 µm. **B)** Expression of *T/BRACHYURY (T/BRACH)*, *SOX2*, and *CDX2* in undifferentiated hPSCs and NMPs (n = 3). **C)** Immunofluorescence images showing T/BRACH and SOX2 colocalization in hESC-derived (upper panels) and hiPSC-derived (lower panels) NMPs. Scale bar: 50 µm. **D)** Quantification of T/BRACH and SOX2 signal colocalization using Pearson’s correlation coefficient (n = 3).

Building on studies showing that NMPs self-organize into neuromuscular organoids (NMOs) [4,21], we incorporated the same progenitor population into a defined fibrous architecture. NMPs were encapsulated using our custom RoWS system, which produces cell-laden hydrogel fiber with controlled core-shell compartmentalization [8,9,10](**Fig. 3**). The system supports high-throughput fiber production, with a typical extrusion rate of approximately 4.2 m/min and a bioink deposition rate of approximately 0.5 mL/min, and collects the fibers into aligned bundles. NMPs were suspended in a fibrinogen-based core surrounded by an alginate shell; subsequent thrombin treatment formed the fibrin matrix within the core. The core–shell configuration combined a cell-supportive matrix with an external boundary that defined the construct’s initial geometry. During culture for up to 50 days, encapsulated cells proliferated, elongated, and differentiated into muscle and neural compartments within the nmTubs (**Fig. 3**).

**Figure 3.**
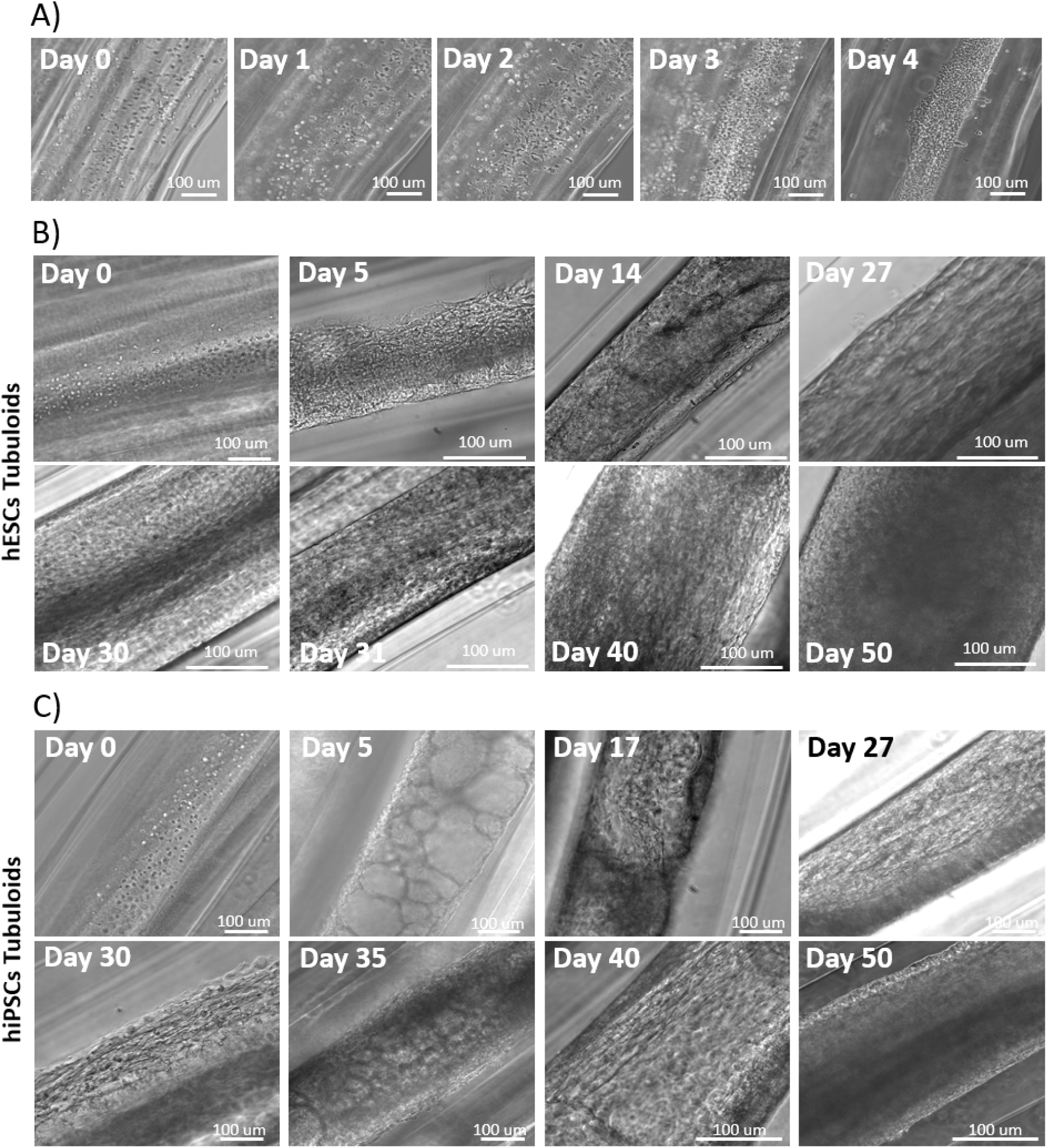
Morphological development of neuromuscular tubuloids (nmTubs) during prolonged culture. **A)** Representative brightfield images of hESC-derived NMPs within hydrogel fibers during the first four days after biofabrication. Day 0 denotes NMPs encapsulation by RoWS and the start of nmTub culture. B, C) Representative brightfield images showing the morphological progression of hESC-derived (**B**) and hiPSC-derived (**C**) nmTubs from day 0 to day 50. Scale bars: 100 µm.

We compared nmTub development from hESC- and hiPSC-derived NMPs under identical culture conditions, using a Neurobasal-based medium initially supplemented with growth factors that support neuro-myogenic differentiation [4,21]. Immediately after biofabrication (day 0), rounded NMPs were distributed along the hydrogel core (**Fig. 3A**). Between days 2 and 4, cells elongated along the fiber axis and formed increasingly dense cellular assemblies. By day 5, hESC-derived nmTubs exhibited a compact, relatively uniform morphology, whereas hiPSC-derived nmTubs contained more visibly distinct compartments (**Fig. 3B, C**). Ki67 immunostaining confirmed substantial proliferative activity at day 5, with positive cells accounting for 77.79% and 71.32% of the population in hESC- and hiPSC-derived nmTubs, respectively (**Fig. S1D, E**). Both constructs underwent further organization and compaction between days 14 and 30. By days 30–50, they displayed thicker cellular structures and greater central optical density in brightfield images (**Fig. 3B, C**). Spontaneous contractions appeared in hESC-derived nmTubs by day 14 and persisted through days 30–50 [VIDEO SV1, SV2], whereas contractile activity in hiPSC-derived nmTubs remained weak [VIDEO SV3]. Morphological examination identified regions resembling developing neural and muscle tissues in both constructs (**Fig. 4A, B**). Neural rosette- and neural tube-like structures were particularly evident in hESC-derived nmTubs, while some hiPSC-derived constructs contained elongated cells with branched projections (**Fig. 4A**). Both cell lines also generated elongated, locally aligned structures resembling myotubes (**Fig. 4B**), consistent with concurrent neural and myogenic differentiation within the fibers. Between days 45 and 50, hESC-derived nmTubs developed organoid-like protrusions that retained spontaneous contractile activity (Fig. S1C; [VIDEO SV4]). These structures likely emerged as continued tissue growth generated internal mechanical stresses that exceeded the resistance of the surrounding alginate shell, causing local rupture and allowing the tissue to expand beyond the original fiber boundaries.

**Figure 4.**
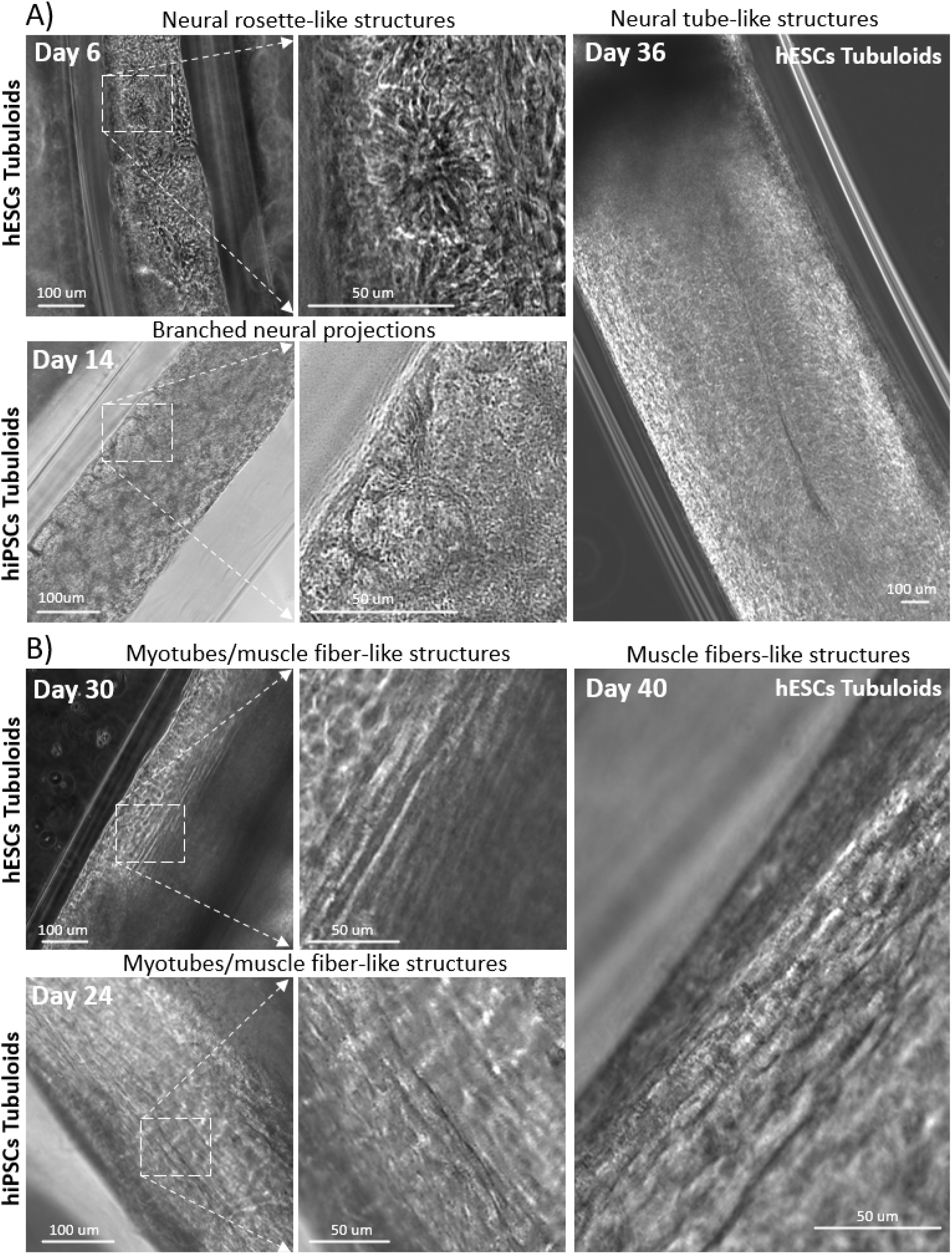
Emergence of neural- and muscle-like structures in neuromuscular tubuloids. **A)** Representative brightfield images showing neural rosette-like structures, branched cellular projections, and neural tube-like structures in hESC- and hiPSC-derived nmTubs. Scale bars: 100 µm. **B)** Representative brightfield images of elongated, locally aligned structures resembling developing myotubes and muscle fibers. Scale bars: 50 µm and 100 µm, as indicated.

### 2.2. Temporal progression *of* neural and myogenic *differentiation*

To compare their suitability for generating a neuromuscular junction (NMJ) model, we cultured hESC- and hiPSC-derived nmTubs under identical conditions and profiled changes in neural and myogenic gene expression throughout development (**Fig. 5**, **Fig. 6**). Analyses covered the NMP stage (day 0), early progenitor differentiation (day 5), initial tissue formation (day 14), and later stages of maturation (days 30–50), allowing us to compare the timing and progression of neuromyogenic differentiation between the two cell lines.

**Figure 5.**
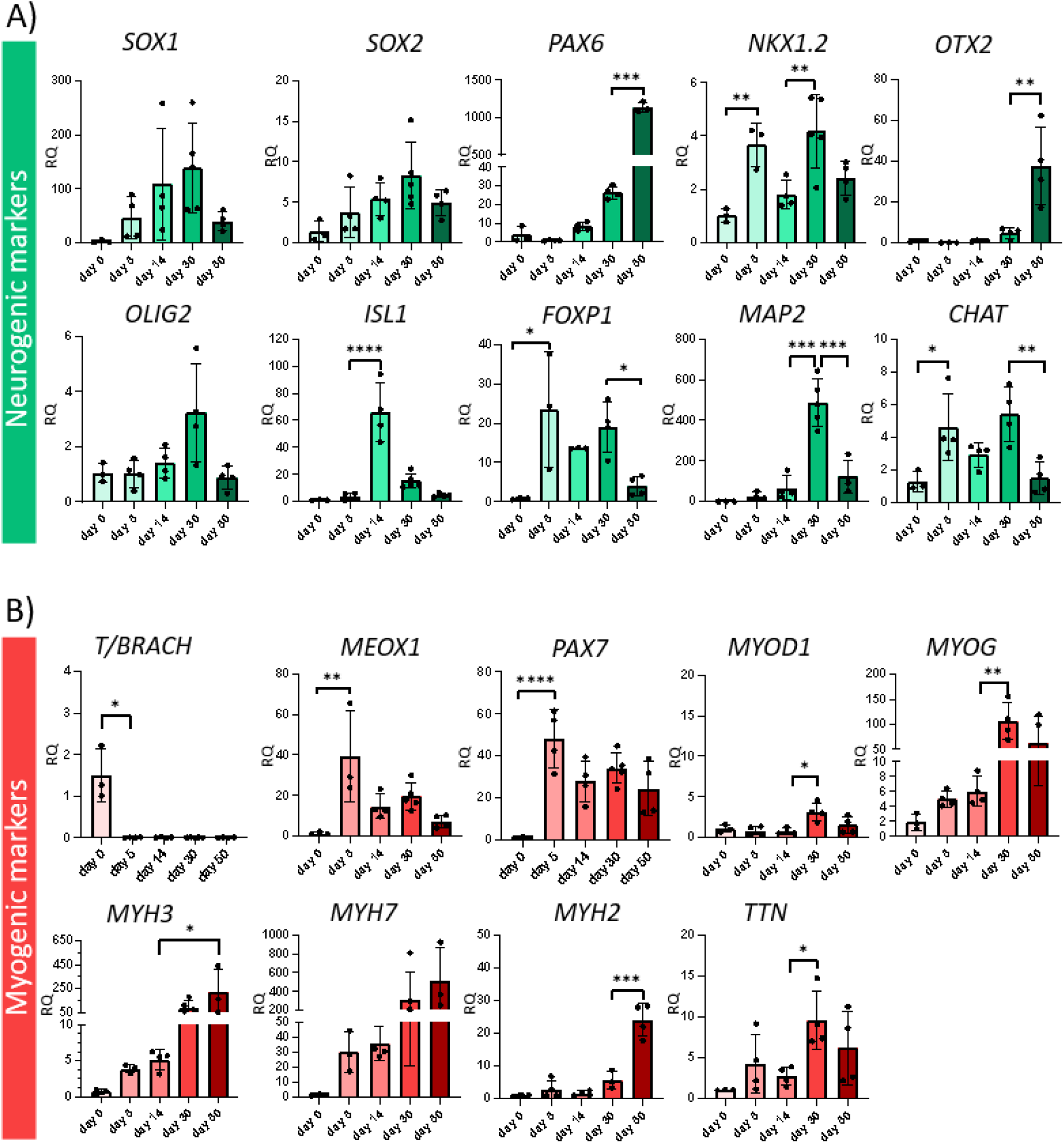
Temporal expression of neural and myogenic markers in hESC-derived nmTubs. A) qPCR analysis of markers associated with neuroectoderm (*SOX1, SOX2*), neural progenitor (*PAX6*), axial identity (*NKX1.2, OTX2*), motor neuron differentiation (*OLIG2, ISL1, FOXP1*), and neuronal maturation or cholinergic identity (*MAP2, CHAT)*. **B)** qPCR analysis of marker associated with mesodermal differentiation (*T/BRACH*, *MEOX1)*, myogenic progenitors and commitment (*PAX7, MYOD1),* and muscle differentiation and maturation (*MYOG, MYH3, MYH7,* MYH2*, TTN)*. *n* ≥ *3. Data are presented as mean ± SD. Statistical significance was assessed by one-way ANOVA: *p* ≤ *0.05, **p* ≤ *0.01, ***p* ≤ *0.001, and ****p* ≤ *0.0001. RQ, relative quantification; qPCR, quantitative polymerase chain reaction*.

**Figure 6.**
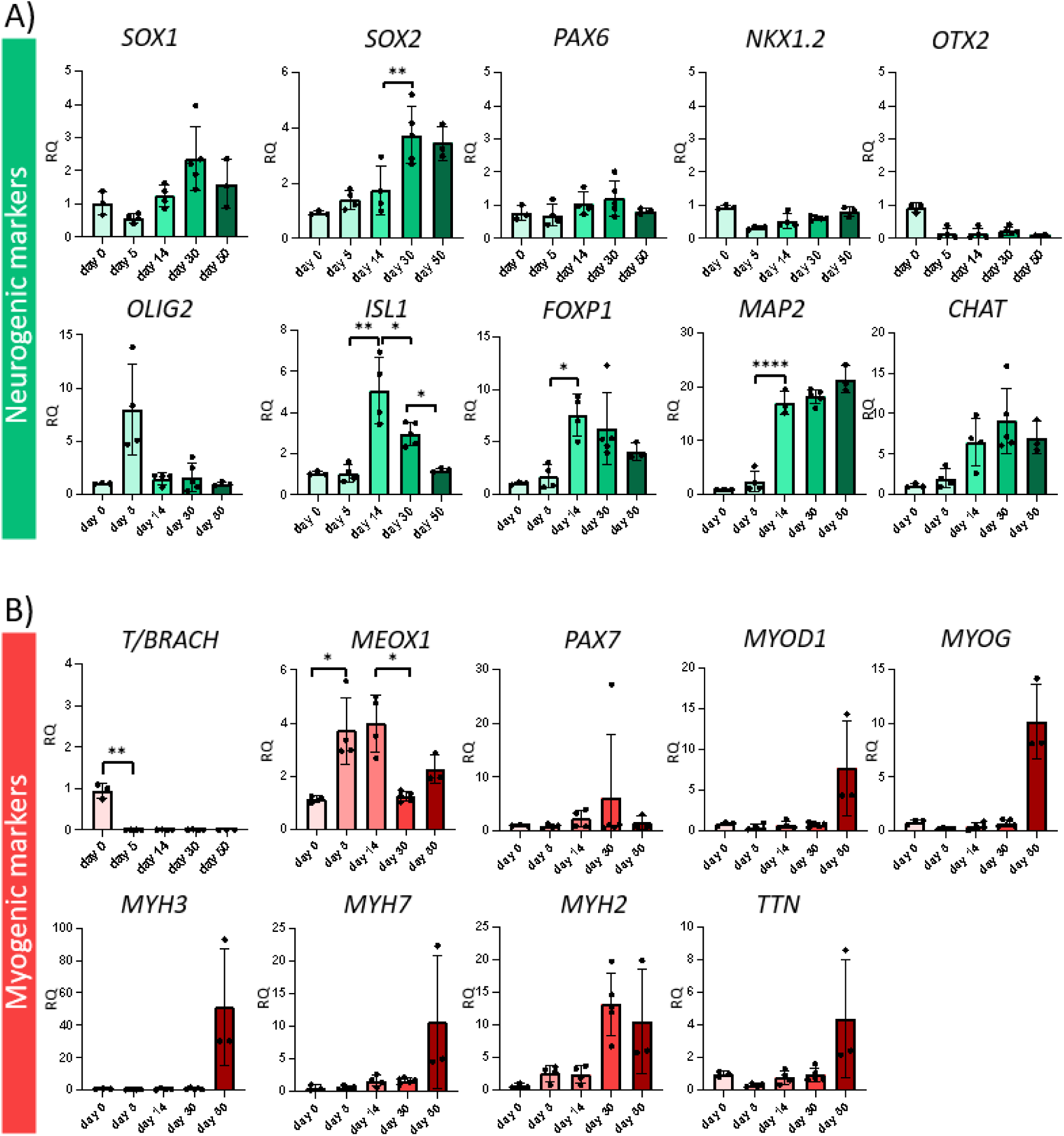
Temporal expression of neural and myogenic markers in hiPSC-derived nmTubs. A) qPCR analysis of markers associated with neuroectoderm (*SOX1, SOX2*), neural progenitor (*PAX6*), axial identity (*NKX1.2, OTX2*), motor neuron differentiation (*OLIG2, ISL1, FOXP1*), and neuronal maturation or cholinergic identity (*MAP2, CHAT)*. **B)** qPCR analysis of marker associated with mesodermal differentiation (*T/BRACH*, *MEOX1)*, myogenic progenitors and commitment (*PAX7, MYOD1),* and muscle differentiation and maturation (*MYOG, MYH3, MYH7,* MYH2*, TTN)*. *n* ≥ *3. Data are presented as mean ± SD. Statistical significance was assessed by one-way ANOVA: *p* ≤ *0.05, **p* ≤ *0.01, ***p* ≤ *0.001, and ****p* ≤ *0.0001. RQ, relative quantification; qPCR, quantitative polymerase chain reaction*.

#### 2.2.1. Progressive neural and myogenic differentiation *in hESC-derived* nmTubs

In hESC-derived nmTubs, *SOX1* and *SOX2* expression increased from day 0 to day 30, consistent with progressive neural differentiation, while *PAX6* expression increased significantly between days 30 and 50 (**Fig. 5A**). During embryonic development, NMPs contribute to neural and mesodermal tissues of the posterior body axis and express *NKX1.2* **[22]**. Their derivatives acquire rostrocaudal positional identities, as reflected in *HOX* gene expression [22,23]. In nmTubs, *NKX1.2* expression increased significantly at days 5 and 30, whereas the anterior marker OTX2 remained low until day 50 (**Fig. 5A**). This profile supports posterior neural differentiation, with the late increase in *OTX2* potentially reflecting greater cellular heterogeneity. *HOXC6, HOXC9,* and *HOXC10*, associated with brachial, thoracic, and caudal thoracic/lumbar identities, respectively [24], were upregulated by day 30 (**Fig. S2A**). *HOXC10* showed a particularly pronounced expression response, consistent with the emergence of posterior positional programs. *OLIG2, ISL1*, and *FOXP1*, associated with motor neuron progenitors and differentiation, increased with distinct temporal patterns between days 5 and 30, while *MAP2* and *CHAT* were strongly upregulated at day 30 (**Fig. 5A**). These findings support progression from neural progenitor states toward neuronal differentiation and a cholinergic phenotype. Glial-associated markers also increased: *SOX10* was significantly upregulated at days 5 and 14, and *GPR37L1*, associated with spinal astrocytic populations [25], remained elevated from days 5 to 50 (**Fig. S2B**). GFAP immunostaining further identified astrocyte-like cells at later stages of development (**Fig. S2C**), indicating that neural differentiation was accompanied by the emergence of supportive glial populations.

Myogenic gene expression also changed progressively during hESC-derived nmTub development (**Fig. 5B**). *T/BRACH* expression was high at the NMP stage and decreased significantly by day 5, while *MEOX1* increased markedly, consistent with progression toward paraxial mesoderm and early myogenic specification. *PAX7* and *MYOD1* increased between days 5 and 30, followed by a significant upregulation of *MYOG* at day 30. Between days 30 and 50, expression increased for *MYH3*, associated with embryonic muscle, and for *MYH7* and *MYH2*, associated with slow- and fast-twitch myosin isoforms, respectively [26] (**Fig. 5B**). *TTN,* encoding the sarcomeric protein titin, was also significantly upregulated. Together, these profiles indicate progressive myogenic differentiation and acquisition of muscle maturation-associated molecular features.

#### 2.2.2. Distinct differentiation kinetics *in hiPSC-derived* nmTubs

hiPSC-derived nmTubs exhibited a distinct temporal pattern of neural gene expression (**Fig. 6A**). *SOX1* and *SOX2* increased gradually, whereas *PAX6* did not increase significantly. Sustained *NKX1.2* expression and low *OTX2* levels were consistent with a posterior neural program. Genes associated with motor neuron differentiation (*OLIG2, ISL1, FOXP1*), neuronal maturation (*MAP2*), and cholinergic identity (*CHAT*) were also expressed. However, the less pronounced *ISL1* and *MAP2* responses, together with the morphological observations, suggested less extensive neural differentiation than in the hESC-derived constructs.

Myogenic differentiation also occurred in hiPSC-derived nmTubs but followed a later temporal profile (**Fig. 6B**). *T/BRACH* expression declined after NMP induction, while *MEOX1* increased at early time points. *PAX7* remained relatively low, with only a modest increase at day 30. In contrast, *MYOD1*, *MYOG*, *MYH3, MYH7, MYH2,* and *TTN* reached their highest expression levels around day 50. These findings indicate delayed acquisition of myogenic differentiation-associated features in the hiPSC line examined and demonstrate that a shared biofabrication and culture protocol can yield distinct developmental kinetics depending on the starting cell population.

### 2.3. Spatial organization and cellular composition *of* nmTubs

We assessed early myogenic organization by examining PAX3-positive progenitors at day 5. PAX3-positive cells formed clusters in both nmTub types (**Fig. 7A**) and accounted for 3770 and 1245 PAX3-positive nuclei in hESC- and hiPSC-derived constructs, respectively (**Fig. 7B**). The observed difference in the early proportion of PAX3-positive progenitors may reflect differences in the dynamics of myogenic progenitor specification and later muscle tissue development between the analyzed cell lines. By day 30, immunostaining revealed spatially distinct muscle and neural domains with different relative proportions (**Fig. 7C, D**). TTN-positive muscle tissue occupied approximately 21867 µm^2^ (7.23% of the area) in hESC-derived nmTubs and 3737 µm^2^ (12.34% of the area) in hiPSC-derived nmTubs; this difference was not statistically significant. In contrast, TUBB3-positive neural tissue occupied 100846 µm^2^ (36.1%) and 2184 µm^2^ (8.15%), respectively, with a significantly greater neural area in hESC-derived constructs (**Fig. 7D**). This difference was consistent with the neural gene expression profiles. In hESC-derived nmTubs, neuronal projections extended toward and closely associated with TTN-positive muscle regions, whereas hiPSC-derived constructs contained more localized neural clusters with less extensive projections. Thus, both cell lines generated muscle and neural compartments within the same fibrous architecture, but hESC-derived nmTubs developed a more extensive neural network for subsequent investigation of neuromuscular interactions.

**Figure 7.**
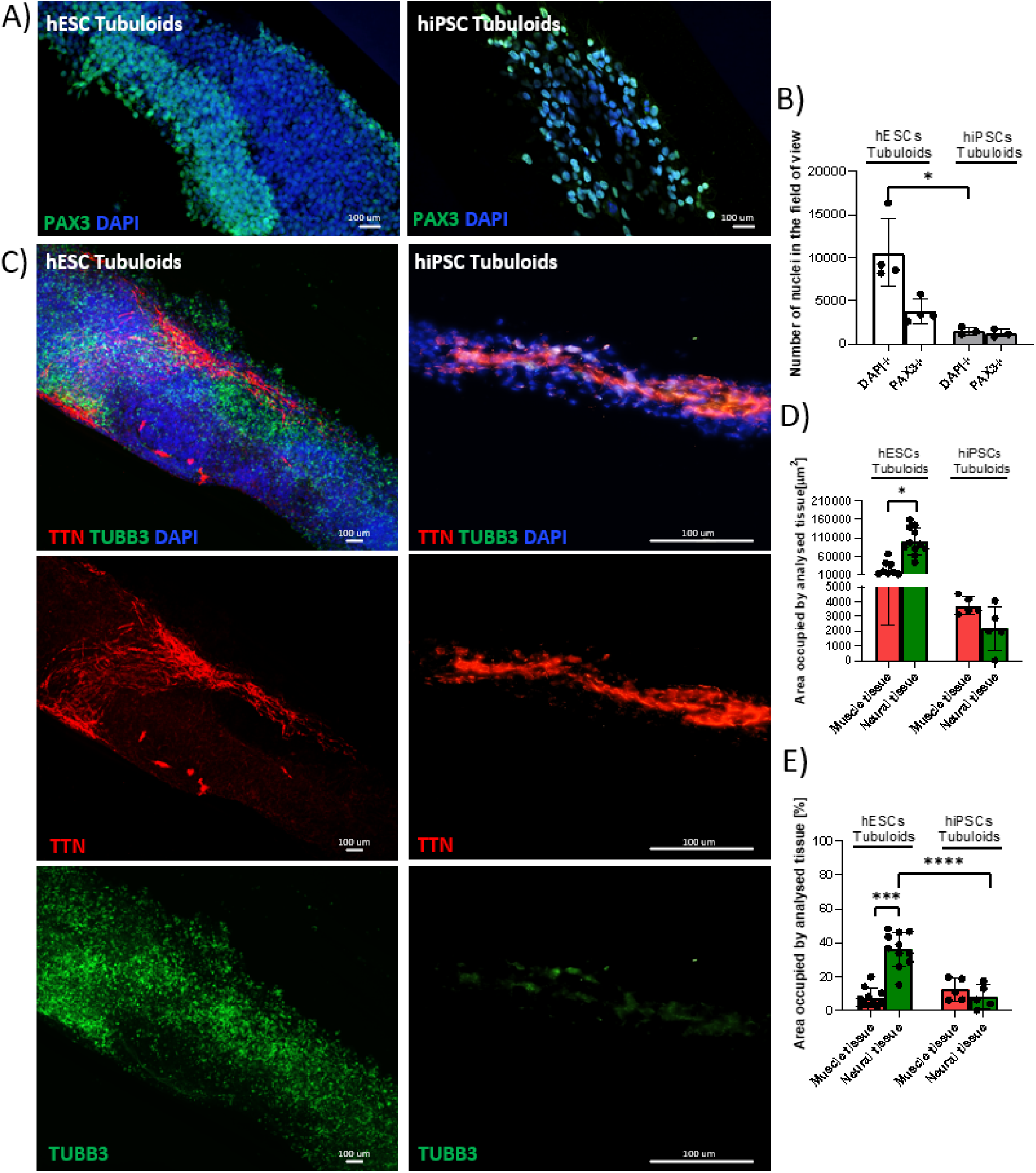
Cellular composition and spatial organization of hESC- and hiPSC-derived nmTubs. **A)** Immunolocalization of PAX3□ myogenic progenitors at day 5. Scale bar: 100 µm. B) Quantification of PAX3□ cells at day 5 (n ≥ 3). C) Immunofluorescence images showing skeletal muscle tissue (TTN, red), neural tissue (TUBB3, green), and nuclei (DAPI, blue) at day 30. Scale bar: 100 µm. **D)** Area occupied by the muscle and neural tissue at day 30 (n≥3). **E)** Percentage of area occupied by muscle and neural tissue at day 30, normalized to the area of th tubuloid in the field of view (n ≥ 3).

Because a subset of PAX3 somitic progenitors contributes to muscle and endothelial cell lineages during embryonic development [27,28,29], we examined endothelial-associated differentiation within nmTubs. Both cell lines expressed *VWF, VEGFA, VEGFB, VEGFR1,* and *VEGFR2*, with generally more pronounced or sustained responses in hESC-derived constructs (**Fig. 8A, B**). Immunostaining identified CD31 endothelial-like cells and capillary-like structures in both nmTub types (**Fig. 8C, D**). These structures were more extensive and elongated in hESC-derived nmTubs, whereas hiPSC-derived constructs contained fewer and less organized CD31-positive regions. CD31-positive endothelial tissue occupied approximately 5226 µm^2^ (4.06% of the area) in hESC-derived nmTubs and 1672 µm^2^ (3.72% of the area) in hiPSC-derived nmTubs (**Fig. 8E, F**). These results indicate endothelial-associated differentiation within the multicellular constructs, although they do not establish the formation of a functional vascular network.

**Figure 8.**
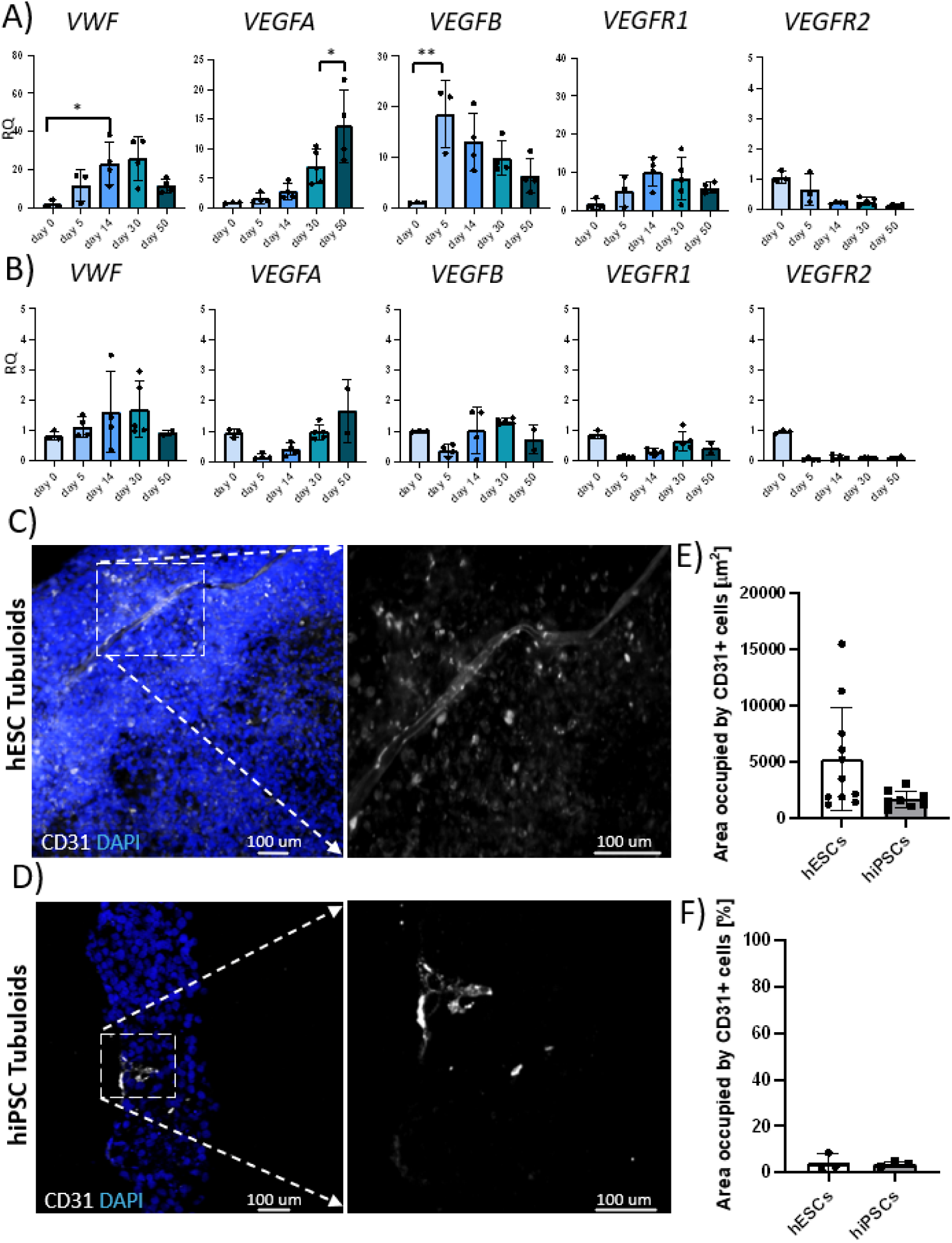
Endothelial-associated differentiation in hESC- and hiPSC-derived nmTubs. A, B) qPCR analysis of VWF, VEGFA, VEGFB, VEGFR1, and VEGFR2 in hESC-derived (**A**) and hiPSC-derived (**B**) nmTubs (n ≥ 3). **C**, **D**) Representative immunofluorescence images showing CD31-positive endothelial-like cells and vessel-like structures (white) in hESC-derived (**C**) and hiPSC-derived (**D**) constructs. Nuclei were counterstained with DAPI (blue). Scale bars: 100 µm. **E)** Area occupied by CD31-positive structures (n=3). **F)** Percentage of area occupied by CD31-positive structures, normalized to the area occupied by nuclei in field of view (n = 3). RQ, relative quantification; qPCR, quantitative polymerase chain reaction.

Overall, hESC-derived nmTubs exhibited more extensive muscle-, neural- and endothelial-associated organization, earlier myogenic differentiation-associated changes, and more pronounced spontaneous contractile activity than the hiPSC-derived constructs examined here. We therefore selected hESC-derived nmTubs for subsequent analyses of tissue maturation, acetylcholine receptor organization, and functional calcium transients.

### 2.4. Structural maturation *and* tissue alignment *in hESC-derived* nmTubs

We compared muscle and neural tissue areas and individual myotube diameters between days 30 and 50 to assess structural maturation (**Fig. 9A, B**). Muscle tissue area increased significantly, while neural tissue area remained relatively stable (**Fig. 9C**). By day 50, the muscle and neural compartments appeared as more clearly delineated domains within the constructs (**Fig. 9A**). TTN-positive myotubes also increased significantly in diameter and showed greater structural continuity, with staining features suggestive of developing sarcomeric organization (**Fig. 9B, D**). TUBB3-positive projections remained closely associated with muscle regions (**Fig. 9A**). These findings indicate continued muscle growth and structural maturation alongside the persistence of an extensive neural compartment during prolonged culture.

**Figure 9.**
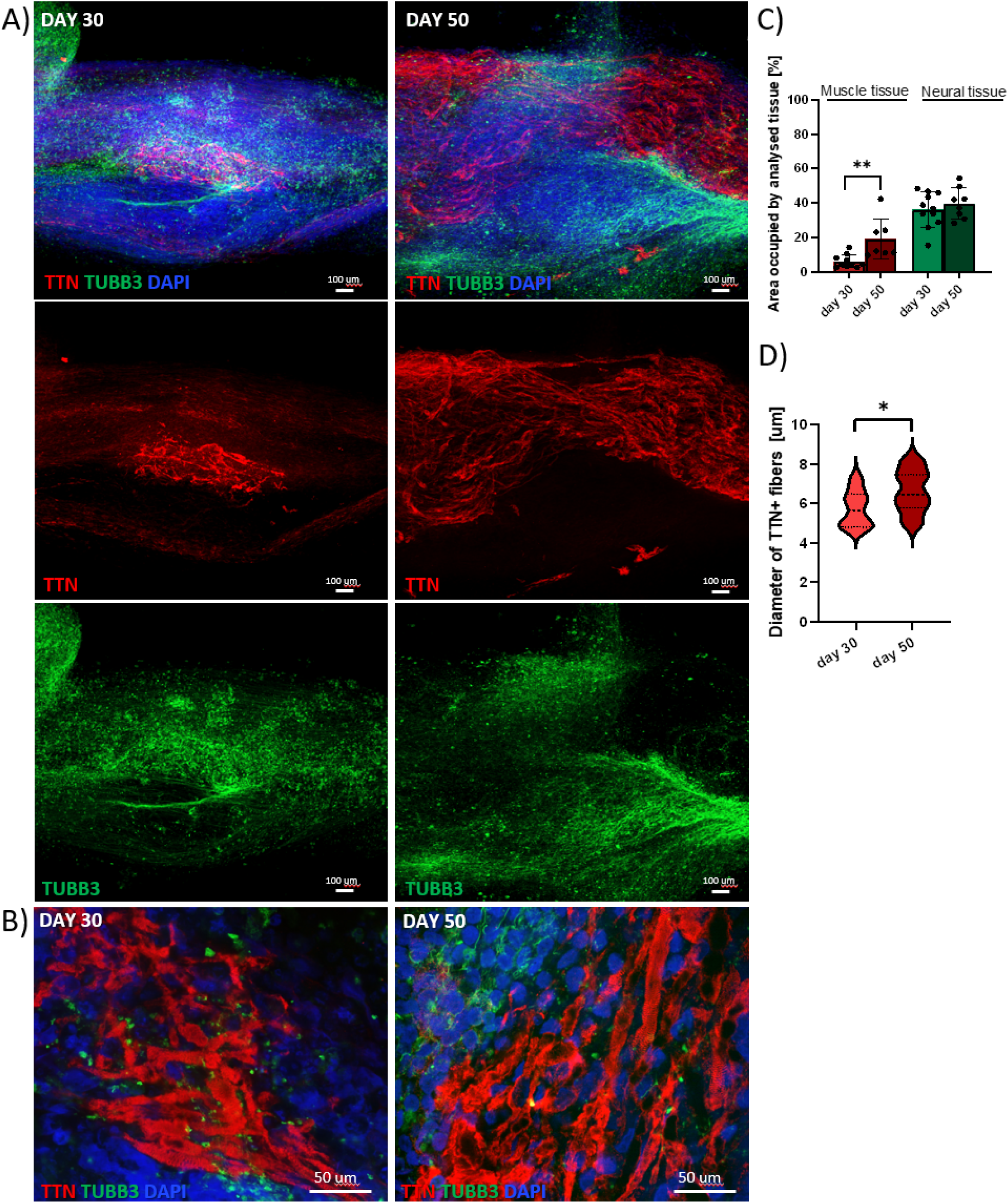
Structural maturation of hESC-derived nmTubs between days 30 and 50. **A)** Immunofluorescence images showing muscle tissue (TTN, red), neural tissue (TUBB3, green), and nuclei (DAPI, blue) at day 30 (left) and day 50 (right). Scale bar: 100 µm. **B)** Representative images showing the spatial association of TTN-positive myotubes and TUBB3-positive neurons at days 30 and 50. Nuclei were stained with DAPI. Scale bar: 50 µm. **C)** Quantification of TTN□ and TUBB3□ tissue areas during maturation (n ≥ 7). **D)** Quantification of TTN-positive myotube diameter (n ≥5).

We quantified the orientation of muscle and neural tissue to examine their organization within the fibrous geometry (**Fig. 10A–E**). Neural projections exhibited a pronounced longitudinal orientation within nmTubs, with greater alignment than the muscle compartment (**Fig. 10B, D, E**). Muscle tissue showed heterogeneous local myotube orientations, including regions of anisotropic organization within the elongated constructs (**Fig. 10A, C, E**). Overall, the defined fiber geometry and compartmentalization supported a common longitudinal framework, while the two tissues differed in their local organization. These results identify neural organization as a prominent feature of the nmTub architecture and distinguish construct-level geometry from the uniform alignment of individual muscle fibers.

**Figure 10.**
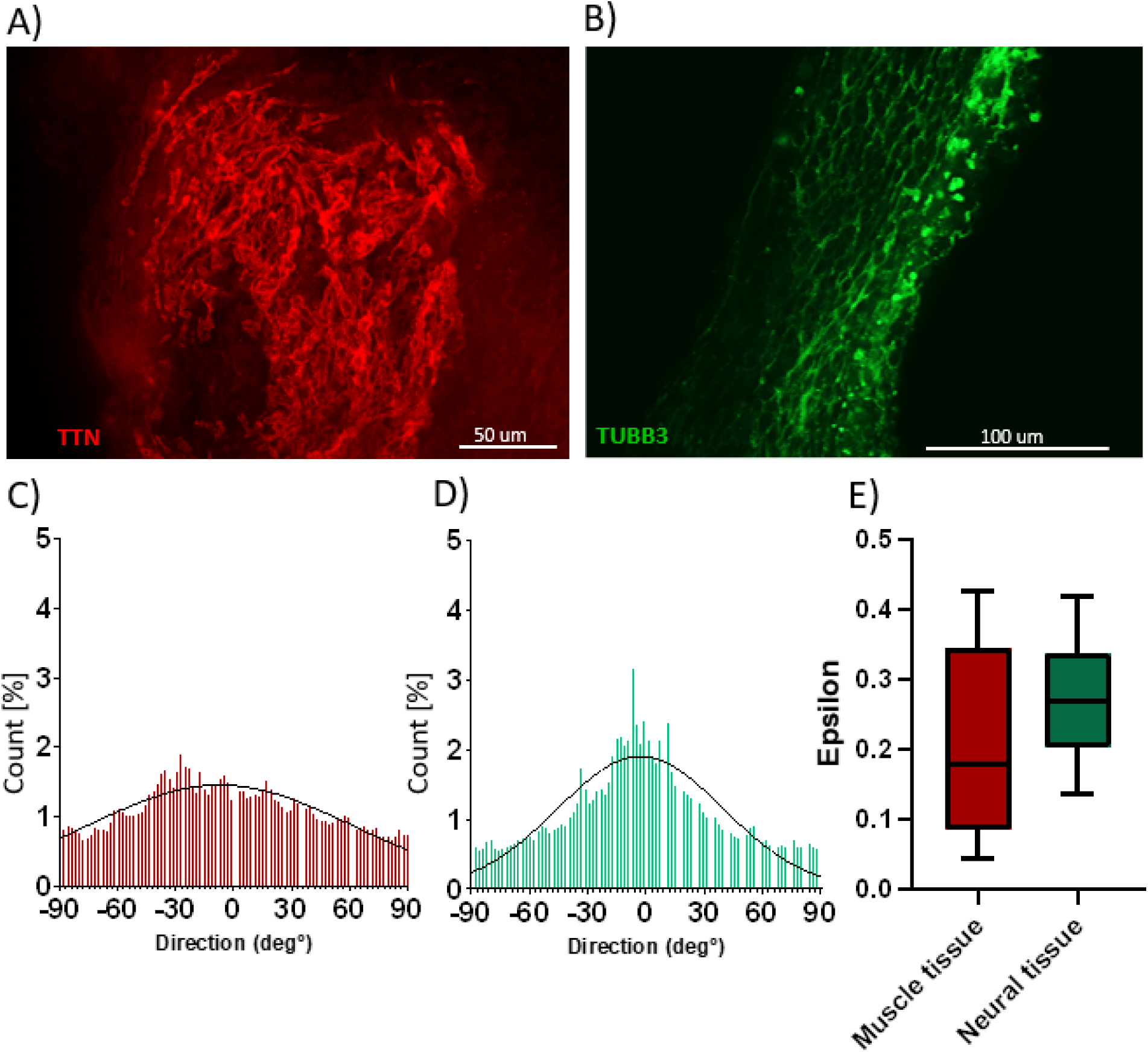
Orientation analysis of muscle and neural tissue in hESC-derived nmTubs. **A, B)** Representative immunofluorescence images of TTN-positive muscle fibers (**A**) and TUBB3-positive neural processes (**B**) at day 30. Scale bars: 50 or 100 µm. **C, D)** Angular intensity distributions used to assess muscle (**C**) and neural (**D**) orientation. **E**) Comparison of the fitted order parameter magnitude, |ε|, between muscle and nervous tissue present in nmTubs (n ≥ 4). Higher values indicate greater uniaxial alignment.

### 2.5. Acetylcholine receptor organization during nmTub *maturation*

At the neuromuscular junction (NMJ), acetylcholine (ACh) released from motor neuron terminals binds to acetylcholine receptors (AChRs) clustered on the postsynaptic membrane of the muscle, triggering depolarization and ultimately muscle contraction [30]. To investigate the emergence of postsynaptic features within nmTubs, we examined AChR distribution using fluorescent α-bungarotoxin (BTX) at days 14, 30, and 50 (**Fig. 11**). AChR clusters were detectable in TTN□ muscle regions by day 14, coinciding with the onset of spontaneous contractile activity (**Fig. 11A; VIDEO SV5**). At this stage, titin staining appeared as discrete puncta rather than organized striations, consistent with early muscle structural development [31]. By day 30, AChR labeling formed relatively large, diffuse domains along TTN-positive fibers **(VIDEO SV6)**. At day 50, these domains appeared smaller and more sharply delineated, suggesting progressive remodeling of receptor organization during tissue maturation (**Fig. 11B, VIDEO SV7**).

**Figure 11.**
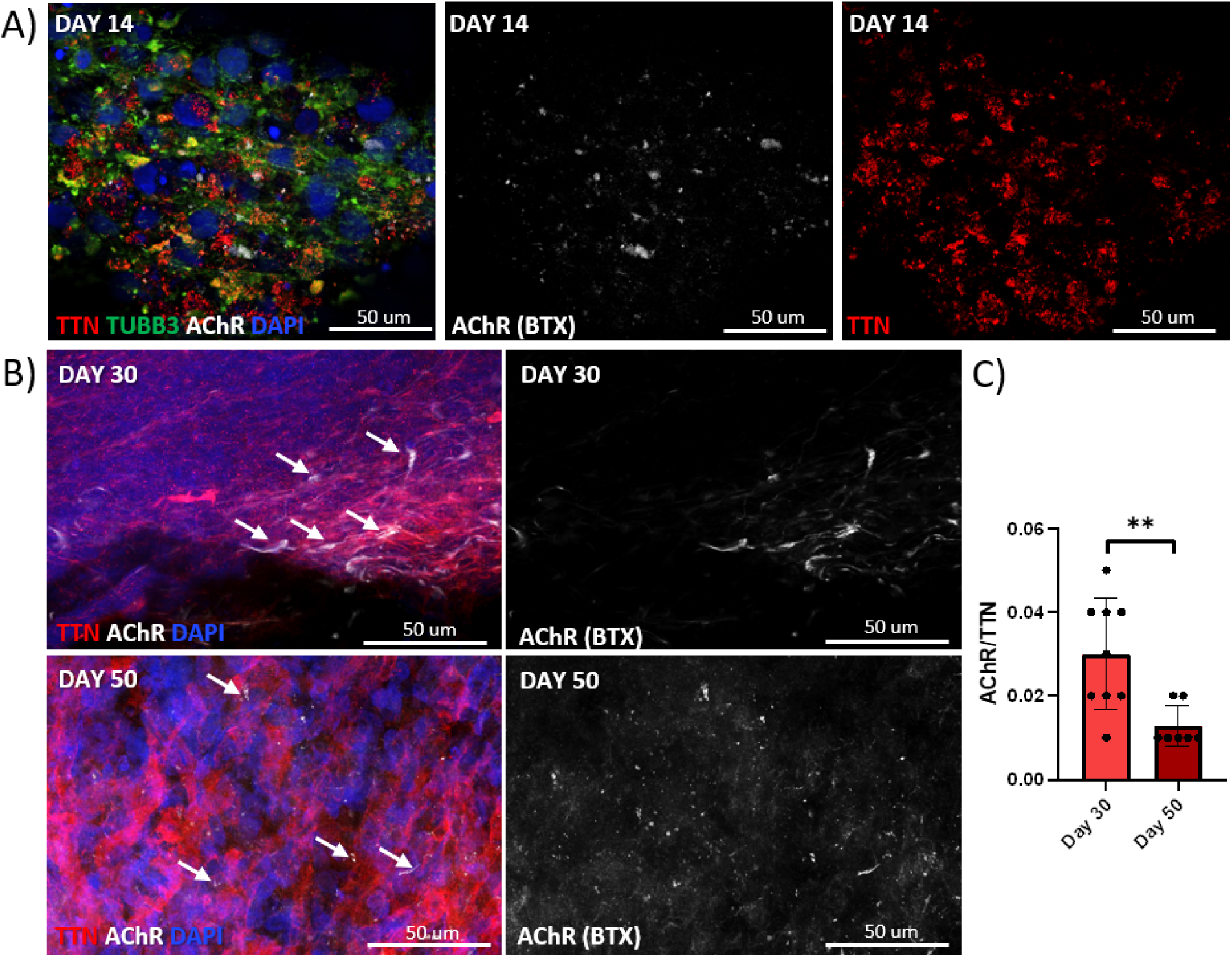
Acetylcholine receptor organization in hESC-derived nmTubs at days 14, 30, and 50. **A)** Immunofluorescence images showing AChR clusters (white), muscle tissue (TTN, red), neural tissue (TUBB3, green), and nuclei (DAPI, blue) at day 14. Scale bar: 50 µm. **B)** Changes in the morphology and distribution of AChR clusters (white, marked with arrow) relative to TTN-positive muscle tissue (red) between days 30 and 50. Nuclei were stained with DAPI (blue). Scale bar: 50 µm. **C)** AChRxy6 area normalized to TTN□ muscle area at days 30 and 50 (n ≥ 7). AChRs were labeled using fluorescent α-bungarotoxin.

Quantification showed a decrease in the AChR□ /TTN□ area ratio between days 30 and 50 (**Fig. 11C**). Together with the morphological changes, this finding is consistent with a more spatially restricted distribution of AChR-rich domains. However, muscle area also increased over this period, so the reduction in the normalized ratio cannot be attributed exclusively to receptor cluster condensation. These observations support temporal remodeling of postsynaptic features but do not, by themselves, demonstrate functional neuromuscular transmission.

### 2.6. Acetylcholine-associated calcium transients *in hESC-derived* nmTubs

Calcium signaling contributes to myoblast fusion, myotube formation, and muscle contraction [32,33,34], as well as neural progenitor proliferation and differentiation [35,36]. To complement the morphological assessment of postsynaptic AChR organization with a functional readout, we monitored intracellular Ca² dynamics by live imaging following ACh stimulation (**Fig. 12A**).On day 14, in some nmTubs we were able to detect cells presenting Ca^2+^ oscillations after ACh stimulation, indicating a responsive cellular population (VIDEO SV8). Quantitative analysis of Ca^2+^transients on day 50 (VIDEO SV9) revealed a trend toward an increased fraction of responding cells and higher relative fluorescence intensity changes (Δ*F/F_min_*) following ACh stimulation compared to non-stimulated (SV 10) constructs. However, these changes were not statistically significant (**Fig. 12B**). While high variability between biological replicates was evident, the persistence of ACh-induced transients at later stages provide evidence of ongoing physiological responsiveness within the constructs. Together with structural maturation, this temporal pattern may reflect a shift in the cell populations exhibiting stimulus-associated activity during nmTub development. Further experiments will be needed to identify the cell populations that respond and determine whether muscle activation occurs directly in response to ACh or via neuronal signaling.

**Figure 12.**
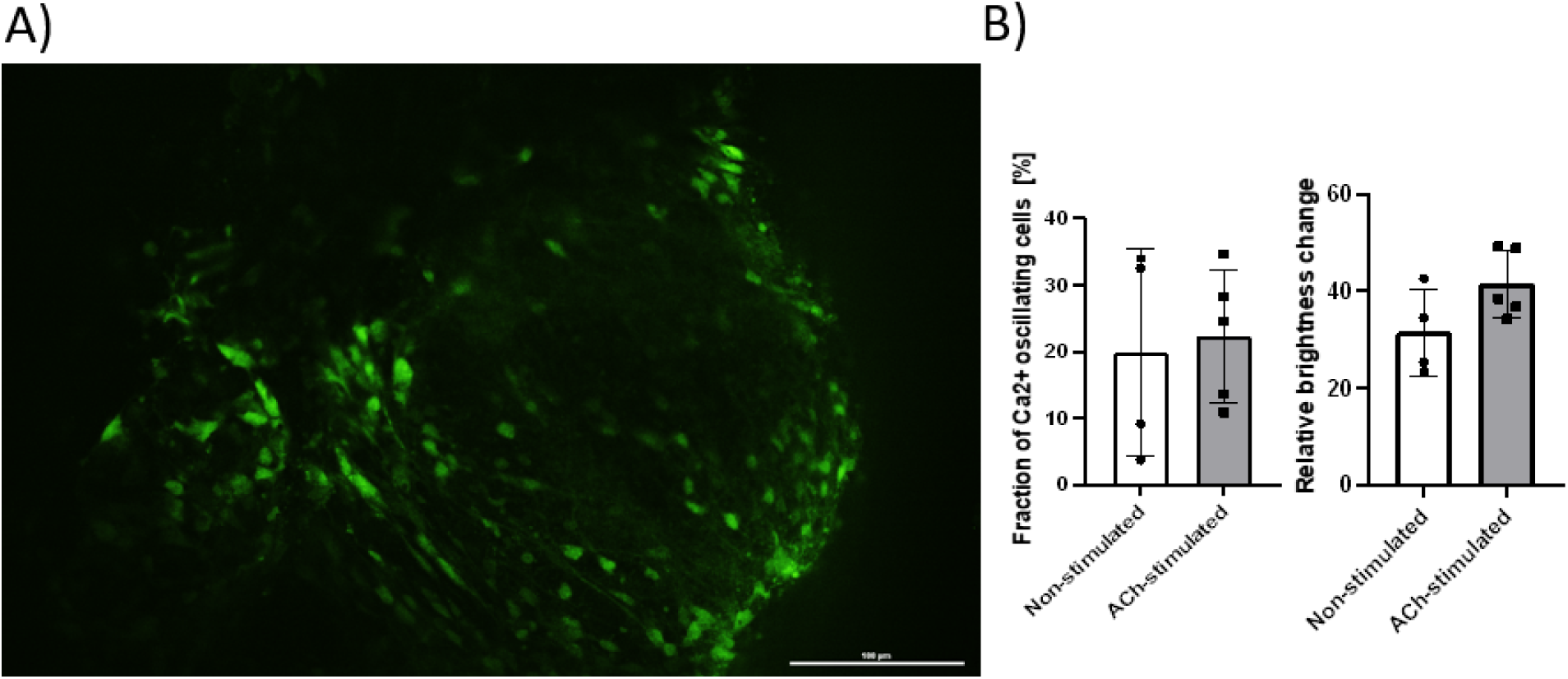
Calcium activity in hESC-derived nmTubs assessed by Fluo-4 imaging. **(A)** Representative fluorescence image of nmTubs stained with the calcium-sensitive dye Fluo-4 on day 50. Bright green fluorescence identifies regions displaying intracellular Ca²□ activity within construct. Scale bar = 100 μm. **(B)** Quantification of spontaneous calcium oscillations in non-stimulated and acetylcholine (ACh)-stimulated nmTubs. Left panel shows the fraction of Ca²□-oscillating cells, while the right panel presents the relative fluorescence intensity chang (n ≥ 4).

## 3. Discussion

Reconstructing human neuromuscular tissue requires both cellular complexity and spatial organization. Skeletal muscle contains a hierarchical arrangement of myofibers, extracellular matrix, and supporting populations, including satellite cells, fibroblasts, motor neurons, Schwann cells, and vascular cells [1,2]. Existing two-dimensional cultures, microphysiological systems, organoids, and bioprinted tissues reproduce different aspects of this organization. Our approach combines a developmentally competent progenitor population with a biofabricated core-shell architecture, enabling muscle, neural, and additional supporting populations to emerge through self-organization within a geometrically defined, anisotropic environment.

Two-dimensional models remain valuable for accessible, reproducible experiments and high-throughput assays, but they incompletely represent the three-dimensional interactions of native muscle [3]. Microphysiological muscle-on-chip systems address part of this limitation by embedding myogenic cells within hydrogels and applying uniaxial tension through anchoring structures [2]. Such models have supported studies of muscle regeneration [37], neuromuscular junction formation and neuromuscular disease [6,38], and contractile function [39]. Extending these systems to larger or more heterogeneous tissues remains challenging because construct geometry, mechanical loading, and cellular composition must be coordinated [1,40]. nmTubs offer a complementary approach in which a common progenitor population generates multiple tissue compartments within a continuous fibrous matrix.

Biofabrication provides tools for the precise 3D placement of cells and biomaterials within engineered muscle tissues [41,42]. Extrusion-based strategies have enabled the assembly of myogenic, neural, and tendon-associated compartments into defined architectures [10,42]. Furthermore, our previous studies demonstrated that muscle constructs biofabricated via wet spinning can promote functional muscle regeneration in models of volumetric muscle loss [8].

Nevertheless, reproducing the fine structure of native tissue remains difficult, and incorporating separately differentiated cell populations introduces additional requirements for cell sourcing and culture compatibility [43]. These requirements become more complex as supporting populations are added [44]. Here, NMPs provide a common developmental starting point for neural and mesodermal differentiation [14,17,45], while RoWS establishes a defined core–shell architecture that guides tissue development within an anisotropic environment. The fibrin-based core provides a permissive matrix for cellular organization, whereas the alginate shell defines the initial fiber boundary and imposes geometric confinement. Together, these features allow muscle and neural populations to develop within the same construct from a shared progenitor pool, without requiring their separate differentiation and subsequent assembly. Previous NMP-derived neuromuscular organoids [4] and two-dimensional NMJ models [13] established the capacity of this progenitor population to generate interacting muscle and neural tissues. nmTubs extend this developmental strategy into an elongated, anisotropic biomaterial environment, enabling the generation of tissue constructs at substantially larger dimensions than those typically achieved in organoid, 2D, and organ-on-chip models. Unlike organoids, which rely predominantly on intrinsic morphogenesis, nmTubs combine developmental self-organization with the geometric guidance of an anisotropic core-shell scaffold [5]. Within this environment, neural projections exhibited pronounced longitudinal organization, whereas muscle tissue formed elongated domains with heterogeneous local fiber orientations. These findings highlight the platform’s capacity to guide overall tissue architecture, while indicating that uniform alignment of individual muscle fibers requires further optimization.

Beyond architectural guidance, the starting cell population influenced the progression and outcome of tissue development. Under identical biofabrication and culture conditions, hESC- and hiPSC-derived nmTubs exhibited distinct differentiation kinetics and cellular compositions, with the hESC-derived constructs developing more extensive neural and endothelial-associated compartments and more pronounced spontaneous contractile activity. These differences are consistent with previously reported variation in the differentiation potential of pluripotent stem cell lines [46,47]. However, as only one hESC and one hiPSC line were examined, these findings cannot establish broader differences between embryonic and induced pluripotent stem cells or implicate a specific reprogramming mechanism. Validation across additional lines will therefore be needed to assess the consistency of nmTub development and distinguish line-dependent outcomes from features reproducibly supported by the platform.

The expression profiles support progression through developmental stages within the fibrous constructs. High *T/BRACH* and *CDX2* expression at the NMP stage was followed by upregulation of *MEOX1*, consistent with a transition toward the paraxial mesoderm. This sequence agrees with reported links between T/BRACH activity and *MEOX1* transcription [48] and with MEOX1’s role in somitic development [49]. Subsequent changes in neural, glial, and myogenic markers indicate the emergence of several differentiation programs within the same constructs. Expression of *NKX1.2* and *HOXC6*, *HOXC9*, and *HOXC10* further supports posterior neural identities. The detection of *HOXC6* differs from the predominantly thoracic/lumbar profile described in some NMP-derived organoids [4], although bulk gene expression alone cannot establish the abundance or identity of individual motor neuron subtypes. Rostrocaudal specification depends on temporally regulated morphogen and transcriptional programs [50,51] and may also respond to the local signaling environment [52] and matrix geometry [53]. A fibrous format could alter transport distances and cellular exposure to soluble cues, but nutrient transport and signaling gradients were not measured here. Reports that nanofiber geometry and fibrous substrates influence neurite extension and neuronal maturation [53] provide a rationale for testing these mechanisms directly in future nmTub studies.

Spontaneous contractions, AChR clusters, and Ca^2+^ transients were detectable in hESC-derived nmTubs by day 14. This timing is earlier than the day-30–50 window described for AChR clustering and functional NMJ development in neuromuscular organoids [4,15] and the day-30–40 range reported in neuromusculoskeletal organoid models [21]. The early emergence of spontaneous contractions and AChR clusters in nmTubs suggests that geometric confinement within an anisotropic fibrous environment promotes neuromuscular tissue maturation. By guiding tissue organization while preserving developmental self-organization, this environment may favor the onset of these features, although its specific contribution remains to be established through direct comparisons under matched culture conditions. During subsequent culture, muscle area and myotube diameter increased, neural projections remained associated with muscle regions, and AChR labeling became more spatially restricted. Such remodeling is consistent with features of postsynaptic development [30,54,55]. However, the declining AChR/TTN area ratio also reflects changes in the muscle compartment and cannot, on its own, establish receptor condensation. Likewise, spontaneous contractions and ACh-associated calcium responses provide evidence of functional tissue activity, laying the groundwork for future studies to establish synaptic transmission between motor neurons and muscle fibers within nmTubs. The longitudinal neural organization observed within nmTubs is consistent with studies showing that anisotropic biomaterials can guide neurite extension and axonal growth [53,56,57], including tubular scaffolds investigated for spinal cord repair [57]. Together, these findings motivate further refinement of fiber geometry and matrix properties to strengthen neuromuscular integration and provide testable links between biomaterial design and tissue function.

## 4. Current challenges and future perspectives

The present findings identify several opportunities to further develop the nmTub platform. In particular, our observations suggest that the initial cell distribution strongly influences subsequent tissue development, with local variations in cell density potentially shaping cell–cell interactions, differentiation, and the relative expansion of neural and muscle populations. Improving the spatial uniformity of NMP loading, together with systematic variation in initial cell density, could therefore enhance reproducibility and provide a means to guide tissue composition.

These considerations are particularly relevant to the platform’s scale: nmTubs are generated as fibers extending over tens of centimeters, whereas neuromesodermal organoids typically develop at the sub-millimeter to millimeter scale [4,21]. Achieving consistent cellular organization across these substantially longer constructs presents distinct engineering challenges and opportunities. Further optimization of core–shell composition, matrix mechanics, and conditioning protocols could promote more uniform muscle alignment and closer integration of neural and muscle compartments. Complementing the current analysis of selected regions with measurements across entire constructs would help assess how local organization translates to larger-scale tissue architecture.

Validation across additional hESC and hiPSC lines will establish the approach’s robustness and clarify how cell-line variability influences developmental outcomes. Although comparisons with organoid data provide useful context, parallel cultures under matched conditions and appropriate geometric controls will be needed to distinguish the contributions of confinement, material properties, and culture conditions. Direct measurements of matrix mechanics and molecular transport would further inform the design of these experiments.

Finally, spontaneous contractions, ACh-associated calcium responses, and endothelial-associated structures provide a foundation for investigating more advanced tissue functions. Future electrophysiological, optogenetic, and pharmacological studies can determine the extent of functional neuromuscular transmission, while assessment of lumen formation and vascular function will clarify the potential of the endothelial compartment to support increasingly complex tissue models.

## 5. Conclusions

RoWS biofabrication enables the generation of human neuromuscular tubuloids by combining NMP differentiation with a defined core–shell hydrogel architecture. This approach supports the emergence of muscle-, neural-, glial-, and endothelial-associated populations within elongated constructs, with progressive muscle maturation, longitudinal neural organization, early spontaneous contractions, and AChR clustering. The platform introduces control over initial tissue geometry while preserving developmental self-organization, providing a complementary format to conventional neuromuscular organoids. Further validation across cell lines and direct tests of synaptic function will support its development as a model for human neuromuscular biology and biomaterial-guided tissue engineering.

## 6. Materials and methods

The transgenic human ESC and iPSC lines were registered with the Polish Ministry of Climate and Environment under registration number 04-23/2022.

### 6.1. Human pluripotent stem cell culture

Human embryonic stem cells (hESCs, H9) and induced pluripotent stem cells (hiPSCs, IIMCBi002-A) were maintained at 37 °C in a humidified atmosphere containing 5% CO□, with daily medium changes. Both lines were confirmed to have normal karyotypes and were free of mycoplasma. Before differentiation, hESCs were cultured in mTeSR1 medium (STEMCELL Technologies) and hiPSCs in Essential 8 medium (GIBCO) on Matrigel (Corning). Cells were passaged every two days.

### 6.2. NMP induction

hPSCs were differentiated into NMPs using a previously described protocol [4]. Cells were maintained for at least two passages, dissociated with Accutase (Life Technologies) at approximately 70% confluence, and plated as single cells on Matrigel-coated 10 mm culture dishes (Life Technologies). NMP induction was performed for three days in Neurobasal-based (NB) medium supplemented with 3 µM CHIR99021 and 40 ng/mL bFGF (both MedChemExpress). NB medium consisted of a 1:1 mixture of Advanced DMEM/F12 (GIBCO), supplemented with 1× N2 (Capricorn Scientific), and Neurobasal medium (GIBCO), supplemented with 1× NCS21 (Capricorn Scientific), 2 mM L-glutamine (GIBCO), 75 µg/mL bovine serum albumin fraction V (Sigma), and 0.1 mM 2-mercaptoethanol (GIBCO).

### 6.3. Bioink preparation

The core–shell formulation combines a fibrinogen-based core with an alginate-based shell. The core provided a matrix precursor for subsequent fibrin formation, while the shell enabled rapid ionic crosslinking during extrusion. The shell contained 2.65% (w/v) low-molecular-weight alginate (Catalog number A18565.36, Thermo Scientific) and 0.35% (w/v) high-molecular-weight alginate (Catalog number 5902409417401, biomus) in 25 mM HEPES (Thermo Scientific). The core contained 1.4% w/v bovine plasma fibrinogen (Catalog number 2000-70, SEQENS) in 25 mM HEPES with 150 mM NaCl (ACROS Organics). Both formulations were filter-sterilized.

### 6.4. Cell preparation for RoWS

NMPs were detached and dissociated into single cells using Accutase, then resuspended in the fibrinogen-based core bioink at a final density of 10–12 × 10 cells/mL.

### 6.5. RoWS biofabrication and nmTub culture

NMP-laden core–shell fibers were produced using the RoWS system described previously [10]. The core and shell formulations were loaded into separate glass syringes (Hamilton) connected to a syringe pump system (Cetoni). The shell and core were extruded at 320 µL/min and 160 µL/min, respectively, into a 0.6 M calcium chloride bath (Sigma-Aldrich), which crosslinked the alginate shell around the cell-laden core. Fibers were collected on a Teflon drum rotating at 60 rpm to form bundles of 15–30 fibers. Bundles were transferred to six-well plates containing thrombin (2 U/ml; Sigma-Aldrich) in 25 mM HEPES and incubated for 30 min at 37 °C and 5% CO□. After thrombin removal, constructs were cultured using conditions adapted from a neuromuscular organoid protocol [4]. During the first four days, NB medium was supplemented with 2 ng/mL IGF1, 2 ng/mL HGF (both Peprotech), 10 ng/mL bFGF, and 50 µM ROCK inhibitor (MedChemExpress). Medium was replaced every two days. From day 4 to day 50, constructs were maintained in NB medium without these added factors at 37 °C and 5% CO□. Samples were collected and imaged on days 0, 5, 14, 30, and 50. Cell pellets and nmTubs intended for RNA analysis were stored at −80 °C.

### 6.6. RNA isolation and quantitative gene expression analysis

Undifferentiated hPSCs and NMPs were lysed in QIAzol (Qiagen) using a syringe. nmTubs were washed in 25 mM HEPES and homogenized using a syringe or a mortar and pestle with liquid nitrogen. RNA was isolated using the miRNeasy Mini Kit (217004, Qiagen), eluted in nuclease-free water, and treated with the TURBO DNA-free Kit (AM1907, Invitrogen). RNA concentration and purity were measured using a NanoDrop One spectrophotometer (Thermo Scientific). cDNA was synthesized using the High-Capacity cDNA Reverse Transcription Kit (4368814, Invitrogen) according to the manufacturer’s instructions. Gene expression analysis used an RNA input of 1 µg, the primers listed in Table S1, and SYBR Green master mix (2008-1000, A&A Biotechnology). At least three independent experiments were analyzed per time point, with two technical replicates. GAPDH served as the reference gene, and reagent controls were included. Amplification curves were analyzed using LightCycler software (Roche) to determine Ct. 2–ΔΔCT analysis was performed as described by Livak and Schmittgen [58].

### 6.7. Immunofluorescence staining of NMPs and whole nmTubs

Before RoWS, NMP monolayers were fixed in 4% paraformaldehyde (PFA) in PBS for 10 min at room temperature and permeabilized with 0.5% Triton X-100 (Sigma-Aldrich) in PBS for 5 min. Samples were blocked in 4% BSA (Capricorn) in PBS for 1 h, then stained with primary and secondary antibodies as described below. Whole nmTubs were fixed in 4% PFA (Agar Scientific) for 1–2 h at 4 °C, depending on the maturation stage, and incubated for 2 h in 4% BSA and 0.3% Triton X-100 in 25 mM HEPES. Primary antibodies (Table S2) were diluted in the same blocking solution and applied overnight at 4°C with gentle agitation. Samples were transferred to six-well plates and washed in 25 mM HEPES. Secondary antibodies conjugated to Alexa Fluor 488, 594, or 647 (Thermo Fisher Scientific) were diluted 1:200 in blocking solution and applied with DAPI (Invitrogen) for 2 h at room temperature. After washing in 25 mM HEPES, samples were mounted in DAKO mounting medium (Agilent) and imaged using a CrestOptics spinning-disk confocal system mounted on a Nikon microscope with NIS-Elements AR 6.20.02 software. At least three independent experiments were performed for each analysis.

### 6.8. Cryosectioning and section immunofluorescence

At day 30, hiPSC-derived nmTubs were fixed in 4% PFA in 25 mM HEPES for 3 h at room temperature, washed three times in 25 mM HEPES, and incubated overnight in 30% sucrose. Samples were embedded in a solution containing 15% gelatin and 10% sucrose at 42 °C. After solidification at 4 °C, blocks were frozen in isopentane and stored at −80 °C. Sections (10–15 µm) were prepared using a Microm HM 505N cryostat (Microm International) and collected on Superfrost slides (Thermo Scientific). Gelatin was removed by incubation in PBS at 42 °C for 20 min. Sections were blocked and permeabilized in PBS containing 3% BSA and 0.3% Triton X-100 (Sigma-Aldrich) for 1 h at room temperature, then incubated overnight at 4°C with primary antibodies (Table S2) diluted in PBS containing 4% BSA. The following day, sections were washed in PBS and incubated with secondary antibodies as described for whole nmTubs before imaging.

### 6.9. Calcium imaging

Intracellular Ca² dynamics were assessed using Fluo-4 AM (F14201, Thermo Fisher Scientific). At days 14 and 50, nmTubs were incubated for 40 min at 37 °C and 5% CO with 2 µM Fluo-4 AM/Pluronic F-127 in prewarmed, high-glucose, phenol-red-free DMEM (Thermo Scientific). Samples were washed once in prewarmed DMEM and stimulated with 200 µM acetylcholine chloride (Sigma-Aldrich). Fluorescence changes were recorded by time-lapse microscopy with 488-nm excitation. Fluo-4 recordings were analyzed with OrganoCa (v0.4.1), an open-source Python tool we developed to automate the ROI-based ΔF/F workflow otherwise performed manually in FIJI [59]. Each recording (.nd2) is first corrected for lateral drift, and individual cells are segmented with Cellpose [60] on the time-averaged and activity images. A cell-free region is used for background subtraction. For each cell, the background-corrected intensity trace is lightly smoothed (Savitzky–Golay filter, 0.7 s), and a baseline F that accounts for photobleaching is fitted across the frames without transients to obtain ΔF/F□. A calcium event is a peak whose amplitude is at least 5 times the cell’s baseline noise (σ), and at least 10 % ΔF/F□, whose prominence is at least 15 % ΔF/F□ and at least 3σ, and that is at least 2 s from the next event. Cells with a noisy baseline, cells touching the image border, and cells at background level were excluded. Cells with at least one event were counted as responding, and cells were ranked by signal-to-noise ratio (largest event amplitude / σ). For group comparisons, the response amplitude was expressed as ΔF/F_min, as in the manual protocol, and each recording was treated as one independent sample (Welch’s t-test and Mann–Whitney U test). OrganoCa also estimates the false-event rate of each recording by running the complete analysis on inverted traces of the non-responding cells. The source code is freely available at https://github.com/saeedrafieyan/OrganoCa.

### 6.10. Quantification of marker-positive cells

Ki67 and PAX3 were analyzed at day 5, and HB9 at day 30, in hESC- and hiPSC-derived nmTubs. The spot-detection tool in NIS-Elements AR 6.20.02 (Nikon) was used to threshold the marker and DAPI channels. The number of Ki67+, PAX3+, or HB9+ positive cells was expressed as a percentage of the total number of nuclei within each field of view.

### 6.11. Quantification of tissue and AChR-positive areas

Immunofluorescence images were acquired from randomly selected fields and analyzed in NIS-Elements AR 6.20.02 (Nikon). TTN□, TUBB3□, or CD31□ regions were manually delineated, and their areas were expressed relative to the total nmTub area. Analyses at days 30 or 50 included minimum 3 biological replicates, with at least three randomly selected fields of view. BTX-positive regions were used to quantify AChR-associated staining, and their area was normalized to the TTN-positive muscle area. This analysis included at least three independent biological replicates and at least four randomly selected fields of view.

### 6.12. Myotube diameter

**Myotube diameter** was measured in Fiji/ImageJ [59] after spatial calibration using a 100 µm scale bar from an image acquired at the same magnification. Three measurements were made at randomly selected positions along each myotube using the straight-line tool, perpendicular to the longitudinal axis. At least three biological replicates and three fields of view were analyzed.

### 6.13. Orientation analysis

Images were processed in Fiji/ImageJ. Because constructs could curve within a field of view, each image was rotated to identify a relatively straight, well-oriented region, which was selected using the polygon tool. Directionality was measured within this region rather than across the full image. Preprocessing consisted of FFT Bandpass Filter, conversion to 8-bit, and Enhance Contrast with histogram equalization. The Directionality plugin was then applied to the selected region. Angular distributions were exported and plotted in Prism (GraphPad).

### 6.14. Orientation order parameter

The Directionality plugin generated normalized angular-intensity distributions, sampled every 2° from −90° to 90° [61]. Each distribution was fitted to a circular orientation distribution function (CODF) [62]: y(θ) = A · (1 − ε²)/(1 + ε² − 2ε · cos 2θ), where ε describes orientational order and A scales the distribution amplitude. Both parameters were estimated by nonlinear least-squares regression in Microsoft Excel using the Solver add-in and GRG Nonlinear algorithm. Goodness of fit was assessed using R². The magnitude |ε| was used for comparisons: values near 0 indicate isotropic organization, and values approaching 1 indicate strong uniaxial alignment. The sign of ε distinguishes two perpendicular peak orientations relative to the chosen reference axis.

### 6.15. Statistical analysis

Experiments included at least three independent biological replicates unless stated otherwise. Normality was assessed using the Shapiro–Wilk test, and equality of variances using Fisher’s F-test. Two-group comparisons used an unpaired, two-tailed Student’s t-test for normally distributed data or the Mann–Whitney U test when parametric assumptions were not met. Comparisons involving more than two groups used one-way ANOVA followed by Tukey’s multiple-comparisons test for parametric data or the Kruskal–Wallis test for nonparametric data. Results are presented as mean ± SD. Statistical significance was defined as p < 0.05 and indicated by *p < 0.05, **p < 0.01, ***p < 0.001, and ****p < 0.0001.

## Funding

This work was supported by the National Science Centre Poland (NCN) within SONATA BIS 12 project no. 2022/46/E/ST8/00284 to M.C.

## CRediT authorship contribution statement

Conceptualization: MC, AF. Methodology: MC, AF. Investigation: AF, SA, FM, MPiletskaya, MPajouhande, SR, AS. Resources: AF, SA. Software: SR. Formal analysis: AF, SA. Writing – original draft: AF, MC. Writing – review and editing: AF, MC, SA, with input from all authors. Supervision: AF, MC. Project administration: MC. Funding acquisition: MC.

## Acknowledgments

The authors thank Dr Maciej Cieśla of the International Institute of Molecular Mechanisms and Machines, Polish Academy of Sciences, for providing the H9 hESC line. We also thank Wojciech Staniszewski for training and technical support in quantitative fluorescence image analysis using NIS-Elements AR 6.20.02.

## Declaration of generative AI and AI-assisted technologies in the writing process

During preparation of this manuscript, the authors used Microsoft Copilot and ChatGPT to improve language, readability, and organization. The authors are responsible for reviewing and verifying the revised content and for the final publication.

## Declaration of competing interests

The authors declare no conflicts of interest.

## Notes

### Competing Interest Statement

The authors have declared no competing interest.

