## Supplementary files for "RoWS Biofabrication of Neuromuscular Tubuloids from Human Neuromesodermal Progenitors"

**SUPPLEMENTARY FIGURES, TABLES AND VIDEO INFORMATION**

**Figure S1. Proliferation analysis and gene expression profiling of markers associated with pluripotency state**. **A)** qPCR analysis demonstrating dynamic drop in the expression of key pluripotency markers (*POU5F1, NANOG)* during hESCs (**A**) or hiPSCs (**B**) differentiation to NMPs (day 0) and then further differentiation of NMPs from day 0 to 50 in hESC-derived (**A**) or hiPSC-derived (**B**) tubuloids, (n≥3); **C)** Representative brightfield images showing the ability of hESC-derived tubuloids to grow, break the shell, and finally form an organoid-like structure between days 45-50 of in vitro culture. Scale bar 100 µm. **D)** Localization of proliferating cells (Ki67+) on day 5 of hESCs- and hiPSC-derived tubuloids culture. Bar 100 µm; **E)** Quantitative comparison of proliferating (Ki67+) cells in hESCs- and hiPSC-derived tubuloids (n=3); Abbreviations: RQ, relative quantification; qPCR, quantitative polymerase chain reaction.

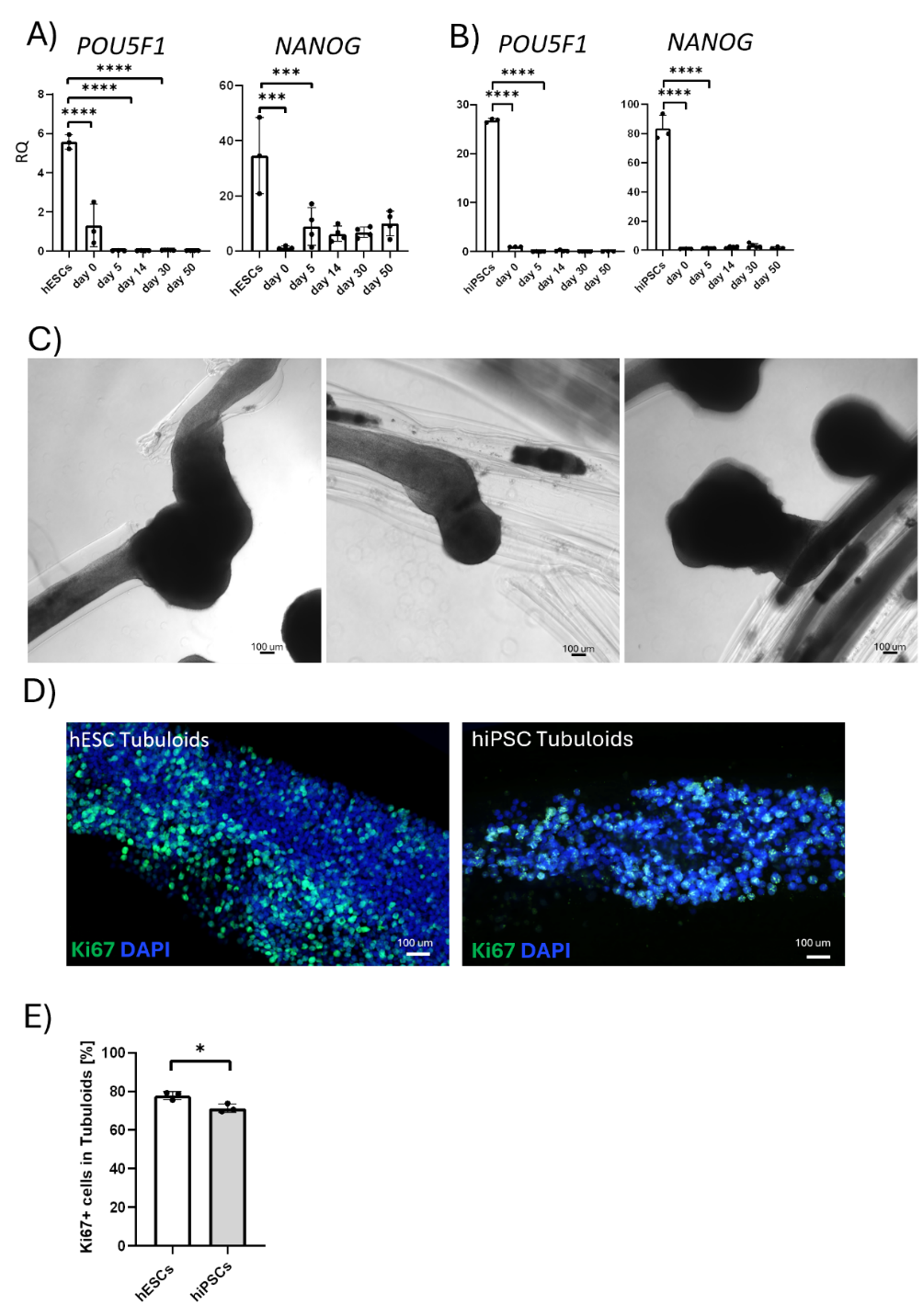

**Figure S2. Gene expression profiling of markers associated with spinal motoneuron subtypes and glia population present in hESC-derived tubuloids**. **A)** qPCR analysis demonstrating expression of brachial (HOXC6), thoracic (HOXC9), and posterior thoracic/lumbar (HOXC10) identity of motoneurons formed in hESC-derived tubuloids, (n≥4); **B)** qPCR analysis demonstrating expression of markers specific for glia progenitors (SOX10) and spinal astrocytes (GPR37L1) formed in hESC-derived tubuloids, (n≥4); **C)** Localization of GFAP+ (white) glia cells in hESC-derived tubuloids on day 30 of in vitro culture. Scale bare 50 um. Abbreviations: RQ, relative quantification; qPCR, quantitative polymerase chain reaction.

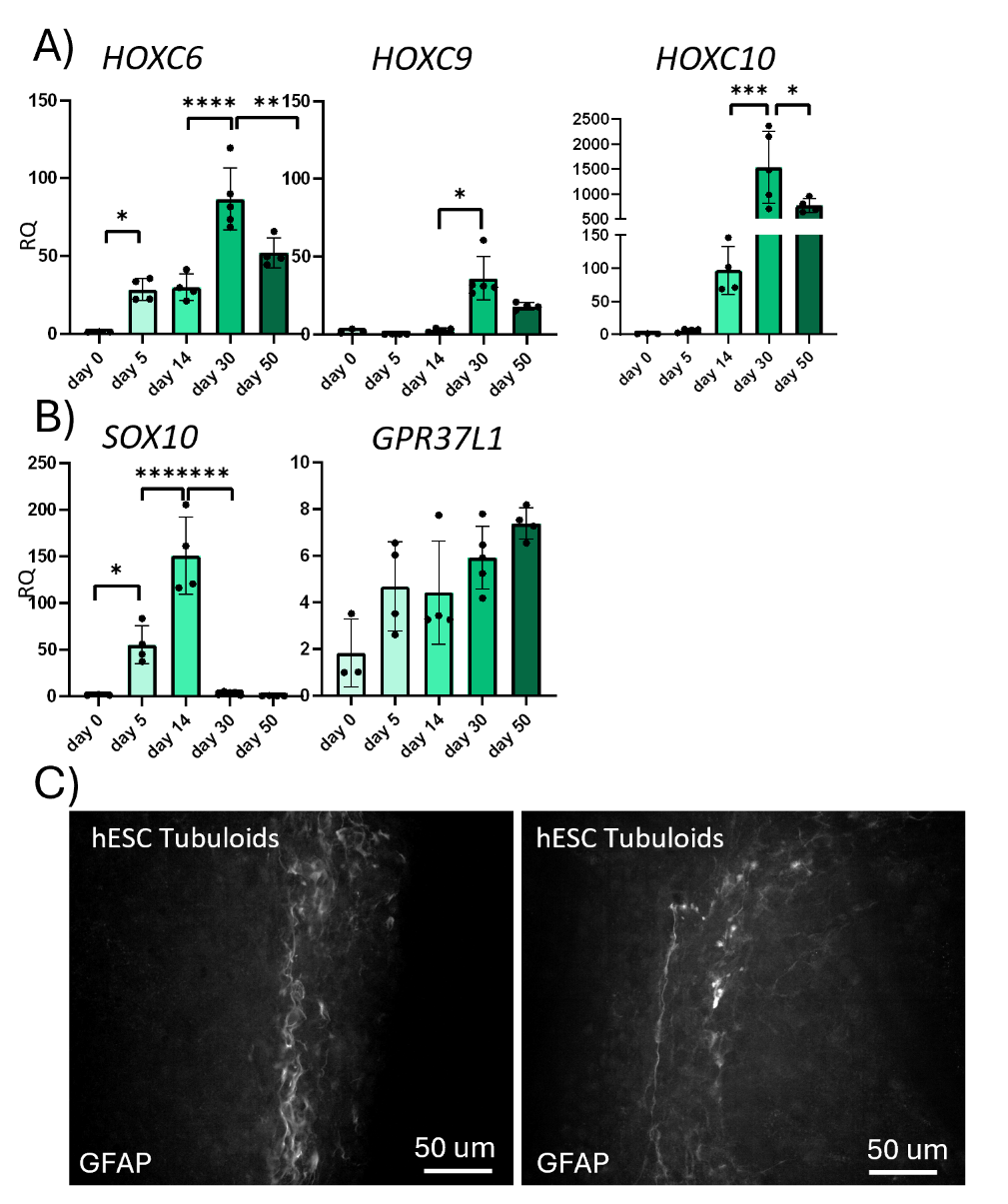

**Figure S3.** **Representative Ca²⁺ transients in hESC-derived nmTubs on day 50 of culture.** Representative graphs of relative fluorescence intensity over time to assess intracellular Ca²⁺ dynamics before (A,B) and after acetylcholine chloride (ACh) stimulation (C, D) in nmTubs. Representative graphs of relative fluorescence intensity, showing the top 16 cells exhibiting Ca²⁺ oscillation before (B) and after (D) ACh stimulation. Analysis was performed in n≥4, for each condition.

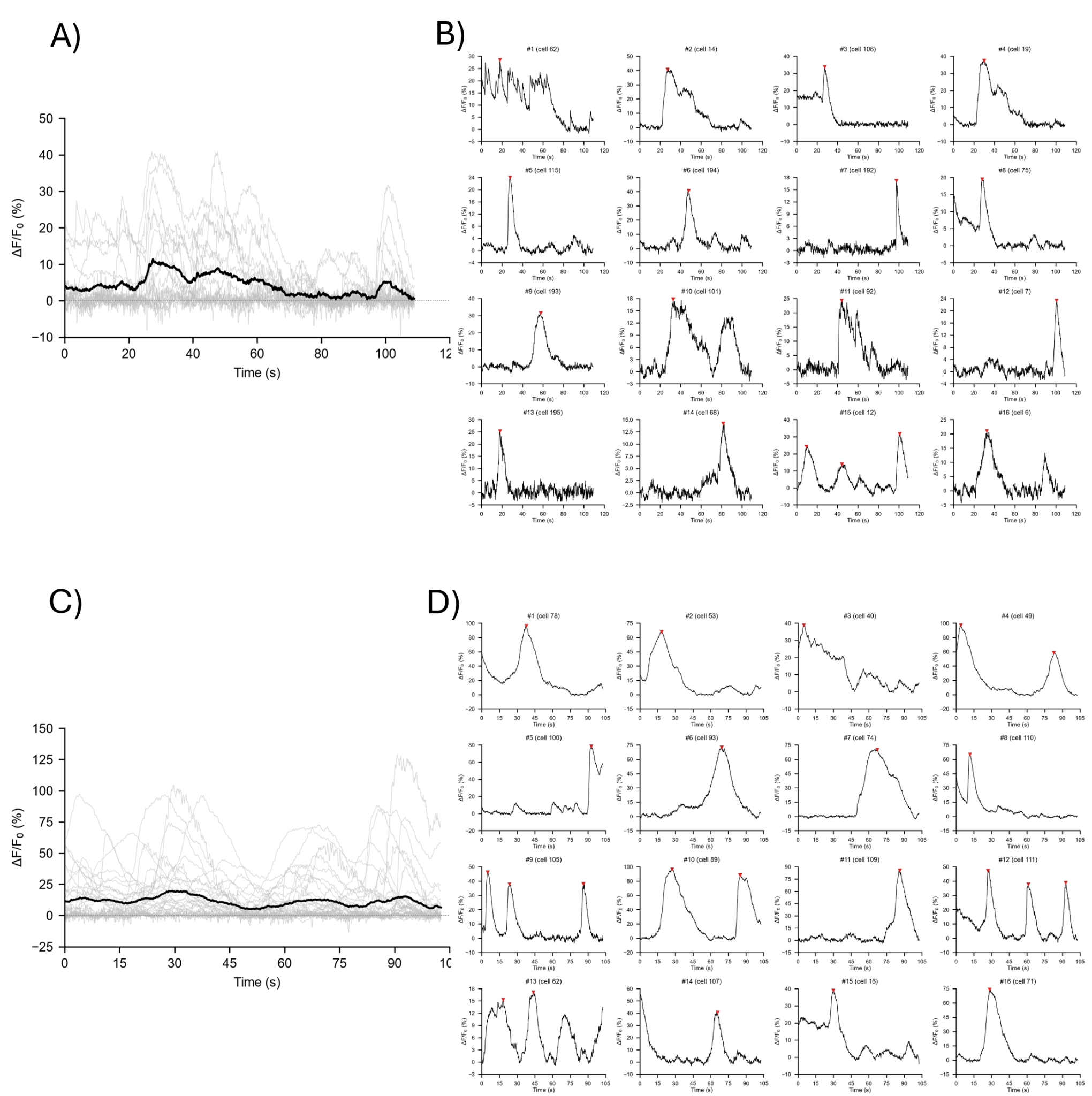

**Table S1. List of primers used in the current study to assess the level of mRNA expression.**

| **Markers** | **Primers** | **Primer sequences** |
| --- | --- | --- |
| NMP | SOX2 | F: GCTACAGCATGATGCAGGACCA  R: TCTGCGAGCTGGTCATGGAGTT |
|  | BRACHYURY | F: TGCTTCCCTGAGACCCAGTT  R: GATCACTTCTTTCCTTTGCATCA |
|  | CDX2 | F: GTCCCAGAGCCCTTGAGTCC  R: GCAGAGTCCACGCTCCTCAT |
| Neurogenic | SOX1 | F: GAGTGGAAGGTCATGTCCGAGG  R: CCTTCTTGAGCAGCGTCTTGGT |
|  | SOX2 | F: GCTACAGCATGATGCAGGACCA  R: TCTGCGAGCTGGTCATGGAGTT |
|  | PAX6 | F: GTACTGAATGACTCAACTGCTCGG  R: CTTTAGAAGGAAGCGACACTCTGC |
|  | OTX2 | F: CAAAGTGAGACCTGCCAAAAAGA  R: TGGACAAGGGATCTGACAGTG |
|  | NKX1.2 | F: AGCTGGTGGCCTTGGAGAAC  R: AGAGACAGCGCGAGGTTCAG |
|  | OLIG2 | F: ATGCACGACCTCAACATCGCCA  R: ACCAGTCGCTTCATCTCCTCCA |
|  | FOXP1 | F: CAAAGAACGCCTGCAAGCCATG  R: TGAGTACTGGCTGAATGACATG |
|  | CHAT | F: TGAGTACTGGCTGAATGACATG  R: AGTACACCAGAGATGAGGCT |
|  | ISL1 | F: AGATTATATCAGGTTGTACGGGA  R: ACACAGCGGAAACACTCGAT |
|  | MAP2 | F: CCCGACGACTCAGCAATGTC  R: TGGCAAGCTGAGGAGATTCG |
| Myogenic | MEOX1 | F: TCTGAGCGCCAGGTCAAAG  R: CTGAACTTGGAGAGGCTGTGG |
|  | PAX7 | F: GGAGGATGAAGCGGACAAGAAG  R: AGGTCAGGTTCCGACTCCACAT |
|  | MYOD1 | F: GAGACCCTCGCAGACCTAAG  R: CTGTAGCACCACACACCATG |
|  | MYOG | F: AAGAAGGGGAGAGGAACAGC  R: GCAACTTCAGCACAGGAGAC |
|  | MYH3 | F: CTCAGGTTGGCTGGAGAAGA  R: AGGGCAGAGACAGTTTGGAA |
|  | MYH7 | F: TCGGAGATGGCAGTCTTTGG  R: CTCCTTCACGGTCACTGTCT |
|  | MYH2a | F: GCTTCTGATTACCACGAACCC  R: CCATAATGCATCACAGCCCC |
|  | TTN | F: TGAGGTGAGCTGGTTTAGGG  R: TTGTCTCACGGTCATGCTCT |
| Pluripotency, spinal motoneuron and glia | POU5F1 | F: CCAGGACATCAAAGCTCTGC  R: CCGCAGCTTACACATGTTCT |
|  | NANOG | F: GTGTTCTCTTCCACCCAGCT  R: CCATTGCTATTCTTCGGCCAG |
|  | HOXC6 | F: TTACCCCTGGATGCAGCGAATG  R: CCGCGTTAGGTAGCGATTGAAG |
|  | HOXC9 | F: CAGCAAGCACAAAGAGGAGAAGG  R: AGTTCCAGCGTCTGGTACTTGG |
|  | HOXC10 | F: GAGCGAAAAGGAGAGGGCCAAA  R: TCCGCTCTTTGCTGTCAGCCAA |
|  | SOX10 | F: ATGAACGCCTTCATGGTGTGGG  R: CGCTTGTCACTTTCGTTCAGCAG |
|  | GPR371L | F: ACGAGATCACCAAGCAGAGGCT  R: GCACAGAGGCTGAAAGTCGTGA |
| Angiogenic | vWF | F: CCTTGAATCCCAGTGACCCTGA  R: GGTTCCGAGATGTCCTCCACAT |
|  | VEGFA | F: TTGCCTTGCTGCTCTACCTCCA  R: GATGGCAGTAGCTGCGCTGATA |
|  | VEGFB | F: AAGGACAGTGCTGTGAAGCCAG  R: TGGAGTGGGATGGGTGATGTCA |
|  | VEGFR1 | F: CCTGCAAGATTCAGGCACCTATG  R: GTTTCGCAGGAGGTATGGTGCT |
|  | VEGFR2 | F: GGAACCTCACTATCCGCAGAGT  R: CCAAGTTCGTCTTTTCCTGGGC |
| Housekeeping | GAPDH | F: CCACATCGCTCAGACACCAT  R: CCAGGCGCCCAATACG |

**Table S2. List of antibodies used in the current study to detect specific cells in tubuloids.**

| **Antibody** | **Host antibody** | **Catalog number** | **Company** |
| --- | --- | --- | --- |
| DAPI | - | D1306 | Invitrogen |
| T/Brachury | Polyclonal goat | AF2085 | R&D systems |
| SOX2 | Monoclonal mouse | MAB2018 | R&D systems |
| Ki67 | Monoclonal rabbit | MA5-14520 | Invitrogen |
| PAX3 | Polyclonal rabbit | ARP32446 | Aviva |
| TTN | Polyclonal rabbit | 27867-1-AP | ProteinTech |
| TUBB3 | Monoclonal mouse, Alexa Fluor 488 conjugate | 53-4510-82 | Invitrogen |
| AChR/BTX | α-Bungarotoxin, Alexa Fluor 647 conjugate | B35450 | Invitrogen |
| GFAP | Monoclonal rat, Alexa Flour 647 conjugate | 51-9792-82 | Invitrogen |
| CD31(PECAM-1) (89C2) | Mouse monoclonal, Alexa Fluor 647 conjugate | 49940S | Cell Signalling Technologies |
| HB9 | Polyclonal Rabbit | PA5-23407 | Invitrogen |

**VIDEO SV1:** Video shows **s**pontaneous contraction in hESC-derived nmTubs on day 14.

**VIDEO SV2:** Video shows **s**pontaneous contraction in hESC-derived nmTubs on day 50. Strong contraction is observed here as compared to hiPSC-derived nmTubs.

**VIDEO SV3:** Video shows **s**pontaneous contraction in hiPSC-derived nmTubs on day 50. The contractions here remained weaker than those of hESC-derived nmTubs.

**VIDEO SV4:** Video depicts organoid like protrusions developed between day 45 and day 50 in hESC-derived nmTubs, the spontaneous contractions were still observed (contraction not shown in video).

**VIDEO SV5:** Video shows the presence of AChR+ clusters in TTN+ muscle regions in hESC-derived nmTubs on day 14. The video was rendered from Z-stack images acquired for slide stained with TTN, BTX and DAPI.

**VIDEO SV6:** Video shows the presence of AChR+ clusters in TTN+ muscle regions in hESC-derived nmTubs on day 30. The clusters are relatively large here along TTN+ fibres**.** The video was rendered from Z-stack images acquired from slides stained with TTN, BTX and DAPI.

**VIDEO SV7:** Video shows the presence of AChR+ clusters in TTN+ muscle regions in hESC-derived nmTubs on day 50. The domains at this stage look smaller and sharply delineated. The video was rendered from Z-stack images acquired from slides stained with TTN, BTX and DAPI.

**VIDEO SV8:** Video depicts presence of calcium ion oscillations in hESC-derived nmTubs on day 14 when stimulated with 200 µM ACh implying a responsive cellular population.

**VIDEO SV9:** Video showing calcium ion oscillations occurring in hESC-derived nmTubs on day 50 in, the oscillations were observed in increased fraction of cells as compared to day 14 of hESC-derived nmTubs on day 50 culture. Stimulation was done with 200 µM Ach.

**VIDEO SV10:** Video shows spontaneous calcium ion oscillations without any stimulation on day 50 of hESC-derived nmTubs culture.
